# LUCID-EV: a robust and quantitative bioluminescent assay for the detection of EV cytosolic delivery in the absence of VSV-G expression

**DOI:** 10.64898/2026.03.24.713260

**Authors:** Louise Merle, Juliette Delas, Lorena Martin-Jaular, Clotilde Thery, Alain Joliot

**Affiliations:** INSERM U932, Institut Curie Centre de Recherche, PSL Research University, F-75005, Paris, France; CurieCoreTech EVs, Institut Curie centre de recherche, PSL-Research University, Paris, France

**Keywords:** cytosolic delivery, extracellular vesicles, bioluminescent complementation assay

## Abstract

Extracellular vesicles (EVs) are key intercellular messengers that modulate the function of target cells by carrying effectors, either at their surface or in their lumen. In the latter case, their action depends on the ability to deliver their content into the cytosol of target cells. How efficiently EVs deliver their content upon interaction with their target cell is thus a central question for understanding the functional impact of this mode of action. To address this question, signal-driven bimolecular interactions between two partners located respectively in the EV lumen and the target cell cytosol have become a widely used strategy to detect the cytosolic delivery EV content. However, the detection of cytosolic delivery with these assays was often tributary to the artificial enhancement of the fusion between EV and cell membranes, usually through VSV-G fusogenic protein expression. Here we provide a robust and quantitative LUCiferase-based complementation assay (HiBiT/LgBiT), to quantify the Internalization and cytosolic Delivery of non-VSV-G-expressing EV content in dendritic cells: LUCID-EV. Most importantly, we show here that the detection of the cytosolic delivery depends on the strategy for loading the HiBiT fragment into EVs and the subcellular location of LgBiT. The cytosolic delivery does not involve the acidic late endosomes environment, mandatory for VSV-G-dependent EV cytosolic delivery. However, the efficacy of cytosolic delivery remains limited and thus, requires highly sensitive assays, suggesting that if physiologically relevant, this might be restricted to specific EV/target cell combinations.

## Introduction

Extracellular vesicles (EVs) are lipid bilayer-enclosed particles secreted by virtually all cell types. They are key effectors of intercellular communication processes, acting in diverse pathophysiological contexts from tissue homeostasis (Buck and Esther N. M. Nolte-‘t Hoen, 2024) to immune system-cancer interplay (Lucotti et al., 2022). Due to their unique and complex structure, EVs can act in different ways. They interact with the surface of target cells, for instance through ligand/receptor interactions (Figure 1A). They can also be taken up by target cells, where their fate and contribution to EV function remain a matter of debate. EVs carry diverse types of biomolecules in their lumen, including proteins and nucleic acids, whose biological activity usually takes place in the cytosol. Among them, mRNAs and miRNAs can accumulate within the EV lumen, and upon EV uptake, they can be translated into proteins in recipient cells (Valadi et al., 2007; Zomer et al., 2015; Rider et al., 2018).

**Figure 1:**
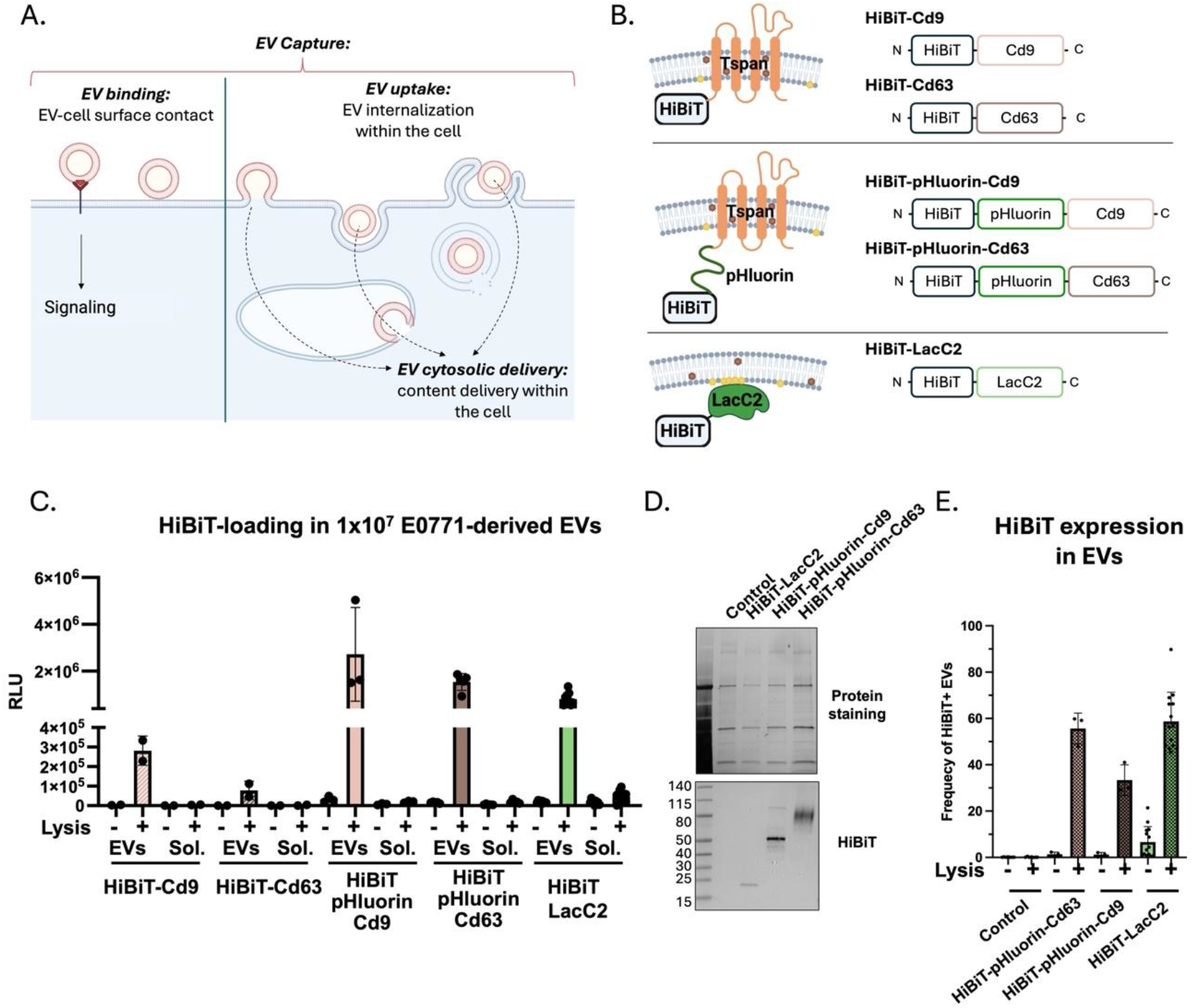
Strategies for efficient HiBiT tag loading into EVs. **A** Proposed modes of action of EVs on recipient cells according to MISEV 2023 guidelines. Created with Biorender. **B**. Description of the fusion protein used to target HiBiT in EVs. **C.** Detection of HiBiT by complementation assay with synthetic LgBiT protein in intact or permeabilized EVs. N= 2 for HiBiT-Cd9, N=2 for HiBiT-Cd63 N= 3 for HiBiT-pHluorin-Cd9, N=4 for HiBiT-pHluorin-Cd63, N=9 for HiBiT-LacC2. **D.** Protein gel staining and western blot detection of HiBiT fusion protein of 5×10^8^ EVs derived from the indicated cell lines). Representative of n=3. **E.** Percentage of HiBiT positive EVs among the total EV population from indicated cell lines quantified by nano-flow cytometry upon or not mild permeabilization (lysis -/+). (Data are expressed as mean ± SEM) N=2 independent experiments for HiBiT-pHluorin-Cd9 and Cd63; N=14 for HiBiT-LacC2

Due to their limited size, EVs can be captured by cells by multiple endocytic pathways (Mulcahy et al., 2014; Gandek et al., 2023; Xiang et al., 2024). A common feature of these processes is that, although topologically inside the cell, EVs remain enclosed in vesicular compartments. It thus requires further steps to deliver their content into the cytosol, involving either membrane fusion events or possibly, membrane rupture (Chahal et al., 2026; Ribovski et al., 2023-6-30). Alternatively, direct fusion of the EV membrane with the plasma membrane could also lead to cytosolic delivery of the EV content (Prada and Meldolesi, 2016; Papareddy et al., 2024). Whatever the mechanism at stake, cytosolic delivery of the EV content needs to be quantified to evaluate its contribution to EV function.

Tracking the intracellular fate of EVs initially relied on fluorescence-based strategies, either by incorporating fluorescent proteins during EV biogenesis (Verweij et al., 2021) or by labelling purified EV components with fluorescent dyes targeting lipid or protein components (Loconte et al., 2023). They proved to be efficient to visualize and quantify EV uptake, but less adapted to demonstrate the cytosolic delivery of the EV content unless combined with a cytosolic reporter in the recipient cell. Following incubation of GFP-Cd63 EVs with HEK293 recipient cells expressing an anti-GFP nanobody fused to mCherry, Joshi et al. observed GFP/mCherry double-positive puncta, consistent with cytosolic cargo release from late endosomes or lysosomes (Joshi et al., 2020). Using a similar strategy based on SunTag peptide and anti-SunTag nanobody expressed in EVs and recipient cells respectively, cytosolic delivery could be observed but only when EVs were loaded with a fusogenic protein (VSV-G) to promote fusion (van den Ende et al., 2026). Cytosolic delivery of EV content was also successfully demonstrated using EV-enclosed luciferase reporter (NanoLucHsp70), although it required additional cell fractionation steps to quantify this process (Bonsergent et al., 2021). Functional cytosolic delivery of EV cargo (protein and RNA) was also elegantly demonstrated using Cre-LoxP- and CRISPR-Cas-based reporter systems by the induction of recombination in recipient cells (de Jong et al., 2020; Liang et al., 2025; Ripoll et al., 2026). Although sensitive, these strategies were less accurate for quantitative assessment of EV cytosolic delivery because the full activation of the reporter in the recipient cell occurs upon a single successful delivery event.

Several strategies based on luminescence bi-molecular complementation have been developed to quantify EV cytosolic delivery in a live cell context. The Split luciferase assay consists in the spontaneous assembly of 2 inactive complementary luciferase fragments, which reconstitute the enzyme activity (Kawamura and Ozawa, 2025). It offers great sensitivity and ease to quantify. The HiBiT/LgBiT split nanoluciferase assay (Dixon et al., 2016) is one of the most sensitive techniques, the two components being respectively located within the donor EV and the cytosol of the target cell, but unfortunately no cytosolic delivery of EV cargo was detected with this assay unless EVs were loaded with a fusogenic protein (VSV-G) to promote fusion (Somiya and Kuroda, 2021). This study aimed to decipher whether optimization of the split nanoluciferase assay allows the detection of cytosolic delivery of EV content in live cells in the absence of VSV-G expression. We focused on three parameters to improve luminescence complementation potential, the loading of the reporter into EVs, the subcellular localization of the cytosolic reporter in the recipient cell and the nature of the EVs and recipient cell type. Here we show that cytosolic delivery of tumor-derived EV content in target dendritic cells is detected, with fast kinetics but limited efficiency. Cytosolic delivery is dramatically increased by VSV-G expression, but through a distinct mechanism from non-VSV-G-bearing EVs. From this study, we demonstrate that optimization of the LgBiT/HiBiT assay allows the detection of EV cytosolic delivery, which might occur through both direct delivery from the plasma membrane and release from endosomal compartments.

## Material and methods

### Cell lines

All cells were kept at 37°C in a humidified atmosphere with 5% CO_2_. The E0771 cell line was cultured in DMEM medium (Gibco) supplemented with 10% sodium pyruvate (Gibco), 10% Fetal Bovine Serum (FBS, Eurobio), 20mM HEPES (Thermo Fisher Scientific), 100U/mL penicillin–streptomycin (Thermo Fisher Scientific), and 2µg/mL puromycin. MDA-MB-231 HiBiT-LacC2 cell line was kindly provided by P. Panagiotis. MDA-MB-231 and HEK-293 FT cell lines were cultured in DMEM medium (Gibco) supplemented with 10% FBS (Eurobio), 100U/mL penicillin–streptomycin (Thermo Fisher Scientific), and 2µg/mL puromycin. The MutuDC cell lines were cultured in IMDM (Sigma-Aldrich), supplemented with 10% FBS (Eurobio), 2mM Glutamax (Gibco), 10mM HEPES, 50μM β-mercaptoethanol, 100U/mL penicillin–streptomycin and 2µg/mL puromycin. All cell lines were regularly tested negative for mycoplasma contamination.

### Lentiviral production and generation of stable cell lines

Lentiviral vectors were all obtained by sub-cloning the fragment of interest into pTwist-Lenti-SSVF (Twist Bioscience). mp-LgBiT and FYVE-LgBiT were obtained by fusion in frame to the N-terminus of the LgBiT coding sequence (CDS) (Dixon et al., 2016) of the myristoylation/palmitoylation (mp) sequence derived from rat lck protein (aa1-14) and a tandem repeat of the FYVE domain of mouse Hrs (aa131-207) respectively. The cytosolic LgBiT correspond to the LgBiT coding sequence (CDS). HiBiT-LacC2 (C2 domain of bovine Mfge8: aa270-427, Uniprot Q95114), HiBiT-Cd9 (mouse Cd9 Uniprot P40240), HiBiT-Cd63 (mouse Cd63 Uniprot P41731), HiBiT-pHluorin-Cd9 and HiBiT-pHluorin-Cd63 were obtained by fusion of the HiBiT sequence (VSGWRLFKKIS) to the N-terminus of the respective CDS. When mentioned, pHLuorin2 CDS (Mahon, 2011) was inserted between the HiBiT tag and the Cd9 or Cd63 CDS. An empty lentiviral vector devoid of any insert was used as control. Lentiviruses were produced by transfection of HEK-293 FT cells with TransIT-293 according to manufacturer instructions with a mix of pCMV-VSV-G (Addgene 8454), pPAX2 (Addgene 12260), and lentiviral vector. Stable transduced cell lines were obtained after 2 weeks selection with puromycin.

### Transient Cell transfection

Transient plasmid transfections were performed on 80% confluent cells using the Lipofectamine 3000 kit according to the manufacturer recommendations. Cells were cultured overnight before medium replacement with fresh culture media for an additional 24h and collection of the conditioned culture media.

### Concentrated conditioned medium and EV isolation by size exclusion chromatography

E0771, MDA-MB-231, MutuDC and HEK-293 cells were grown in complete medium until 80% of confluency. For EV production, conditioned media from 22h culture in serum-free medium (E0771, MDA-MB-231, HEK-293 FT) or medium containing 10% EV-depleted FBS (MutuDC), were cleared of cells and debris by 2 successive centrifugations (300g for 10min and 2000*g* for 20min 4°C) and concentrated on a Centricon Plus-70 Centrifugal Filter (Millipore; MWCO 10kDa) by centrifugation at 2000g at 4°C. When indicated, the concentrated conditioned media (CCM) was loaded on SEC column (Izon qEV single GEN2 35nm) and EVs were collected according to manufacturer instructions. Concentrated CCM and EV aliquots were stored in low protein binding tubes at -80°C. Cells were counted and viability was measured at the end of the culture and always above 85%. EV-depleted serum was obtained by overnight ultracentrifugation of 20% serum-containing medium at 100,000*g* in a 45Ti rotor.

### Nano Flow Cytometry

Particle concentration and size distribution were measured using U30 Flow NanoAnalyzer (NanoFCM) with software version NF profession 2.0. Samples were diluted in 0.22um filtered 10mM HEPES buffer for measurement.

For EV staining, 2×10^8^ SEC purified EVs were incubated with FarRed647-anti-HiBiT monoclonal antibody (1/2000, Promega, CS3278A06, concentration determined after antibody titration with HiBiT and Control EVs) for 20min in ice protected from light, in the presence or absence of 0.2% saponin. Free antibodies were washed away using Exo-spin Mini columns (EXO3, cell guidance systems), following the protocol we recently developed (Nevo et al., 2026).

### Western blot

Cells were lysed with RIPA buffer (NaCl 0.15M, Tris 0.05M, EDTA 2mM, SDS 0.4%, NP40 1%), nuclease (88700, Thermo Scientific) and protease inhibitor (11836170001, Roche) and kept on ice for 30minutes. Cell lysates from (2×105 cells), or 5×108 EVs were resuspended in Laemmli sample buffer (Bio-Rad), boiled 10 min at 95°C, and run in 4–15% Mini-Protean TGX Stain-Free gels (Bio-Rad) in non-reducing conditions. After electrophoresis, proteins were transferred on immuno-blot PVDF membranes using the Trans-Blot Turbo system (Bio-Rad, California, USA). HRP-labelled secondary antibodies were revealed using ECL substrates (1705062, BioRad) for cell lysates, or (34094, Thermo Scientific) for EVs, and imaged with ChemiDoc imager (Bio-Rad, California, USA). The antibodies used were: anti-mouse Cd63 1/200 (clone R5G2, MBL, D263-3), anti-mouse Cd9 1/1000 (clone KMC8, BD Bioscience, 553758), anti-mouse Mfge8 1/1000 (clone 18A2-G10, MBL, D199-3), anti p30 gag MLV 1/1000 (clone R187), env MLV 1/2000 (clone 83A25), anti-HiBiT 1/5000 (N7200, Promega), anti-VSV-G 1/1000 (PA1-30235, Invitrogen).

### LgBiT, HiBiT and DkB production

BL21 DE3 transformed with a plasmid encoding LgBiT CDS fused to a N-terminal his6-bactCherry-Prescission cleavage site tag under the control of T7 promoter (pShCherry2 derivative) were cultured in Magic Media (Invitrogen) overnight at 16°C. The sonicated bacterial lysate was clarified by centrifugation (30.000*g*, 15min) and loaded on hisTrapHP column (Cytivia) according to manufacturer instructions, the his6-bactCherry tag was cleaved by on-column Prescission protease digestion. The protein was eluted and dialysed against phosphate saline buffer (NaCl 100mM, Sodium Phosphate buffer 20mM pH7.5) and protein concentration was adjusted to 50µM and stored in 50% glycerol at -20°C. HiBiT (VSGWRLFKKIS) and DkB (VSGWALFKKIS) peptides were obtained by chemical synthesis (Proteogenix).

### Complementation assay

Cells, SEC fractions or CCM resuspended in 50µL PBS were diluted twice in PBS or 0.05% Passive Lysis Buffer (125mM Tris-HCL pH7.8, 10mM diaminocyclohexaneN,N,N’,N’-tetraacetic acid, 10% glycerol, 1% triton X-100) supplemented with 0.5µM HiBiT (LgBiT complementation) or 0.5µM LgBiT (HiBiT complementation) and incubated for 5 minutes under agitation at room temperature (medium agitation, Spectramax ID3, Molecular Device). Following addition of furimazine (NanoGlo Promega) 1/300 in 50µL PBS and 30 seconds shaking, luminescence was measured for 1 second on Spectramax ID3.

### Proteinase K assay

4×10^8^ SEC-purified EVs were resuspended in PBS and incubated in the absence or presence of 0.2% Triton X-100 and in the absence or presence of 1µg/mL Proteinase K (Invitrogen, PN100005393) for 15minutes at 37°C. Proteinase K was inactivated by the addition of 2 mM cOmplete Protease Inhibitor cocktail (Roche) and subsequent incubation for 15 minutes on ice. Samples were mixed with Laemmli buffer and heated at 97°C for 7 minutes prior to Western blot analysis.

### LUCID-EV assay

1.5×10^5^ MutuDC reporter cells (viability greater than 94%) were seeded in 2 96-well tissue culture-treated white plates (136102, Thermo Scientific) in 50µL CO_2_-independent DMEM (18045088, GIBCO) supplemented with 20µM DkB and incubated at 37°C in a humidified atmosphere with 5% CO_2_. One hour after cell seeding, one of the two plates was transferred on ice (4°C control), and EVs or CCM (adjusted to 10^7^ RLU) diluted in 50µL of CO_2_-independent medium supplemented with 400µg/mL BSA were added to the cells (2 to 4 technical replicates for each condition). An equivalent amount of EVs was used for control condition. Plates were incubated on ice or at 37°C for the specified periods of time. Plates incubated at 37°C were switched to 4°C for five minutes before cytosolic delivery measurement to block intracellular trafficking and ensure comparable measurement conditions between the 4°C and 37°C conditions. Cytosolic delivery was quantified by luminescence after addition of 50µL furimazine (1/300 in PBS) and 2 minutes shaking on Spectramax ID3. To measure uptake, wells were washed twice with 150µL Ca^2+^, Mg^2+^ DPBS (14040117, GIBCO), and cells were lysed in 50µL of 0.5% lysis buffer for 5 minutes at medium shake and luminescence was measured as for cytosolic delivery.

When specified, Bafilomycin A1 (Sigma,100nM final concentration,) or 0.1% DMSO solvent was added to cells 30 minutes before EV addition and in the EV input.

### Anti-HiBiT/IgG treatment of EVs

SEC-purified EVs (adjusted to 10^7^ RLU) were incubated with 0.25µg/mL anti-HiBiT antibody (N7200, Promega) or IgG for 25 minutes at 4°C and subsequently diluted 1/3 before incubation with cells in the LUCID-EV assay.

### Pharmacological compounds affecting endocytosis and intracellular trafficking

MutuDC cells were pretreated with 10µM cytochalasin D (PHZ1063, Thermo Fisher Scientific), 0.3mM L-leucine methyl ester (LLOME; L7393-500MG, Sigma-Aldrich), or 12.5µM Dyngo-4a (HY-13863-MedChem) for 45 minutes at 37°C before the LUCID-EV assay. The compounds were also included during incubation with EVs, which lasted 30 minutes.

### Immunofluorescence

Cells were seeded on Poly-lysine coated wells and cultured for 24 hours in complete medium. Cells were incubated with Wheat Germ Agglutinin (WGA) Alexa Fluor 555 (10 µg/mL, Invitrogen W32464) for 10 minutes at 4°C before fixation with PFA 4% (10 minutes, RT). Cells were incubated with home-made rabbit anti nanoluciferase antibody in PBS 1% BSA 0.5% saponin for 30 minutes protected from light followed by incubation with Alexa647 anti-rabbit antibody (Invitrogen) and mounted in Fluoromont-G with DAPI (Thermo scientific). Images were acquired using Spinning microscope (Inverted Eclipse Ti-E (Nikon) and Spinning disk CSU-X1 (Yokogawa) integrated in Metamorph software. Images were analysed using Fiji software.

### Small EVs and VLPs separation by velocity gradient

EVs and VLPs were separated as described previously (Cocozza et al., 2023). Briefly, conditioned medium cleared of cells and debris by 2 successive centrifugations (300g for 10min and 2000*g* for 20min 4°C) was concentrated up to 6mL on a Centricon Plus-70 Centrifugal Filter (Millipore; MWCO 10kDa) by centrifugation at 2000g at 4°C and centrifuged at 10,000*g* for 16min at 4°C in MLA-80 rotor (Beckman Coulter). The supernatant was ultracentrifuged at 200,000*g* for 50min at 4°C in MLA-80 rotor. The 200k pellet was resuspended in 1mL of PBS and seed on top of a layered 6-22% iodixanol gradient and centrifuged at 187,000*g* for 1h30 at 4°C in Sw32.1Ti rotor (Beckman Coulter). 8 fractions of 2mL were collected carefully from top to bottom, diluted 2-fold in PBS and pelleted at 200,000*g* for 50min at 4°C. Each pellet was resuspended in PBS and stored in aliquots in low protein binding tubes at -80°C.

### LgBiT release assessment

1.5×10^5^ MutuDC reporter cells (viability greater than 94%) were seeded in 96-well tissue culture-treated white plates (136102, Thermo Scientific) in 50µL CO_2_-independent DMEM (18045088, GIBCO) supplemented with 20µM DkB and incubated at 37°C in a humidified atmosphere with 5% CO_2_. Two hours after cell seeding, 50µL of PBS supplemented with 1nM HiBiT and furimazine (1/300) was added to the cells. Release LgBiT in the extracellular space was quantified by luminescence after 30 secondes shaking on Spectramax ID3.

### Statistical analysis

Data were analysed on Excel and statistical analysis were performed on GraphPad Prism 10 with the statistical tests specified in figure legends. Experiments were performed at least twice with different batches of EVs.

## Results

### Selective HiBiT loading in EVs

The HiBiT/LgBiT split nanoluciferase system offers several advantages for cytosolic delivery assays. The affinity of spontaneous interaction between the two fragments is in the nM range, and the small size of the HiBiT fragment (11 aa) is suitable to engineer fusion proteins (Dixon et al., 2016). Our team previously reported that EVs secreted by the murine mammary cancer cell line E0771 are efficiently captured by MutuDC cells and modulate their function (Cocozza et al., 2023). We used this experimental model to monitor EV cytosolic delivery, engineering donor E0771 to secrete HiBiT-loaded EVs and acceptor MutuDC cell lines to express LgBiT reporter. To avoid possible toxicity issues, side effects such as increased EV release (Mathieu et al., 2021) and variability inherent to transient transfection, stable cell lines were generated for each construct. Two strategies were tested to load the HiBiT tag into the lumen of EV (Figure 1B). In the first one, we fused the tag either directly to the N-terminal cytosolic domain of EV transmembrane markers Cd9 or Cd63 tetraspanins (collectively Tspans). Alternatively, the HiBiT tag was spaced with a linker protein (pHLuorin) to move the tag away from the transmembrane domain and enhance its accessibility to the LgBiT fragment. In the second strategy, we fused the HiBiT tag to soluble protein domains that reversibly interact with selected lipids, with the aim of increasing the mobility of the sensor at the membrane interface and thus, its ability to interact with the complementary LgBiT fragment. When expressed in cells, these protein domains interact with the cytosolic face of membrane compartments enriched with the targeted lipid species (Maekawa 2014). The C2 phosphatidylserine-binding domain of Mfge8/Lactadherin (LacC2)(Andersen et al., 2000) (Figure 1B, Supplementary Figure 1A), the mutant D4H cholesterol-binding domain of *Clostridium* Theta toxin (Maekawa and Fairn, 2015) and the PH PIP2-binding domain of PHPLC∂ (Watt et al., 2002) were selected (Supplementary Figure 1A). The three constructs were efficiently expressed in E0771 and addressed to EVs (Supplementary Figure 1B-C). However, HiBiT-LacC2 sensor was detected with a much higher efficiency in the complementation assay with a recombinant LgBiT fragment, both in cell extracts and EVs (Supplementary Figure 1B-C), and thus was kept for the study. HiBiT-LacC2 was then compared to the HiBiT-Tspans as sensors of EV content delivery. All sensors were detected by complementation and western blot in the cell extracts (Supplementary Figure 1D-E), but addition of a linker in the two Tspan fusions significantly increased luminescence activity measured in cell extracts, reaching levels comparable to LacC2 sensor (Supplementary Figure 1D). Next, we quantified the loading of the HiBiT sensors into EVs by measuring luminescence upon incubation with recombinant LgBiT, comparing the EV-enriched (EVs) to the EV-depleted soluble secretome (Sol) separated by SEC. In lysed EVs, all sensors were detected, although with different efficacies, and scarcely with intact EVs due to the restriction of LgBiT access to luminal HiBiT sensors by the EV membrane (Figure 1C). Only residual activity was measured in Soluble fractions (Figure 1C). These data confirmed both the integrity of EV preparations and the efficient targeting of the sensors in the EV lumen. Insertion of a linker protein (pHluorin) between the HiBiT tag and the N-terminus of Tspan sensors increased the luminescence signal up to 20-fold relative to the direct fusion, both in EVs (Figure 1C) and in cell extracts (Supplementary Figure 1D), reflecting improved expression and/or interaction with recombinant LgBiT LacC2 and Tspan-linker constructs were as effective at promoting HiBiT loading into EVs (Figure 1C). We verified by Western blot that they were detected at the expected size in the EV fraction (Figure 1D) and that neither the number of secreted EVs (Supplementary Figure 1F) nor the distribution of Mfge8, Cd9 and Cd63 EV markers (Supplementary Figure 1G) were affected by the expression of the HiBiT sensors. (Supplementary Figure 1F). The percentage of EVs labelled with an anti-HiBiT antibody in mild permeabilization conditions was analysed by nanoflow cytometry: it ranged from 35% (for HiBiT-pHLuorin-Cd9) to 60 % (for HiBiT-Cd63-pHLuorin-Cd63 and HiBiT-LacC2) (Figure 1E, Supplementary Figure 1H-I). As expected, HiBiT detection was significantly decreased in absence of permeabilization. For HiBiT-LacC2 EVs, a minor EV pool (6.6±6.6%) displayed surface-accessible HiBiT (Figure 1E and Supplementary Figure 1H-I). Proteinase K treatment of intact EVs did not result in detectable degradation of the HiBiT-LacC2 sensor, further supporting its main localisation within the EV lumen (Supplementary Figure 2A). Additionally, approximately 70% of LacC2-HiBiT-positive EVs were also positive for at least one of Cd9 or Cd63 tetraspanins (Supplementary Figure 2B). Altogether, these results show that both LacC2 and Tspan-linker sensors are efficiently loaded in the lumen of EVs although a minor LacC2 pool is detected at the surface of EVs.

### Detection of EV cytosolic delivery

Even if delivered into the cytosol of reporter cells, HiBiT sensors would remain associated with membranes, either irreversibly (Tspan sensors) or through lipid interaction (LacC2 sensor). We added a myristoylation/palmitoylation (mp) sequence to the LgBiT reporter to promote the targeting of the fusion protein to the cytosolic leaflet of the membranes of recipient cells in order to increase its probability of interaction with membrane-bound HiBiT sensors. MutuDC cells stably expressing mpLgBiT (MutuDC reporter cells) were generated. We confirmed the expression and complementation of mpLgBiT with a synthetic HiBiT peptide in lysed cell extracts (Figure 2A) and its membrane localization by immunofluorescence (Figure 2B).

**Figure 2:**
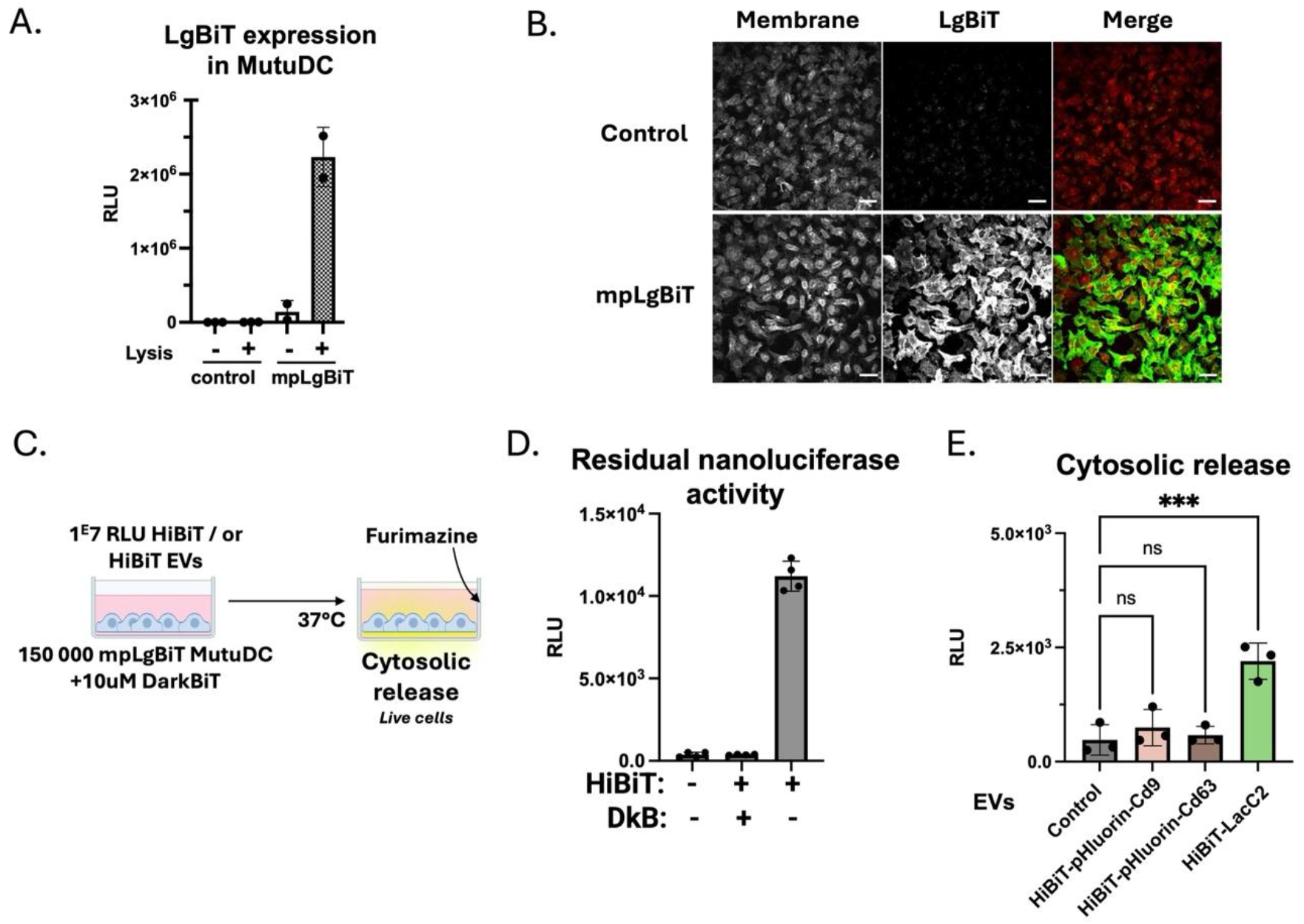
Detection of EV cytosolic delivery by optimization of the complementation assay. **A** LgBiT measurement in reporter cells by complementation assay with synthetic HiBiT peptide. **B.** Spinning disk microscopy images of MutuDC expressing or not mpLgBiT, stained with anti-LgBiT (green) and WGA (plasma membrane, red) and DAPI (nucleus, blue). Representative of N = 2 Scale bar = 10µM **C.** Scheme of the experimental protocol used to measure cytosolic release. Created with Biorender **D.** Measurement of unspecific nanoluciferase activity after MutuDC cells expressing mpLgBiT exposure to HiBiT peptide in presence or absence of DkB peptide **E.** Measurement of luminescence activity following incubation of the indicated EVs with MutuDC expressing mpLgBiT in the presence of DkB peptide. N=3 independent experiments; Error bars = Mean with SD. Statistical analyses were performed using repeated measures non parametric one-way anova with Bonferroni test. \**P* < 0.05, \*\**P* < 0.01, \*\*\**P* < 0.001 and \*\*\*\**P* < 0.0001.

Our assay was designed to minimize experimental manipulations of cells before the measurement of cytosolic delivery. Briefly, MutuDC reporter cells seeded in 96-well plates were incubated in serum-free medium with HiBiT-loaded EVs. At the end of the incubation, the cell-permeable furimazine substrate was added, and luminescence was directly measured after brief shaking (Figure 2C). Due to the extreme sensitivity of the split nanoluciferase assay, residual extracellular luciferase activity coming from cell leakage or cell death was detected after incubation of cells with a cell-impermeable synthetic HiBiT peptide (Figure 2D), thus interfering with the measurement of intracellular cytosolic delivery. As reported by others (Pereira et al., 2019; Solinge et al., 2023), the addition of an excess of an inhibitory peptide (Darkbit, DkB), still able to bind LgBiT but catalytically inactive, was efficient to prevent this contaminating activity when used at 10 µM final concentration (Figure 2D, Supplementary Figure 3A,B). Because the detection of cytosolic release relies on luminescence measurement, the amount of EV input added to the reporter cells was adjusted with the luciferase activity measured in lysed EVs after recombinant LgBiT complementation instead of EV number and set to 10^7^ Relative Luminescence Units (RLU), roughly corresponding to 1500 EVs/recipient cell.

After 2 hours of incubation, no specific activity was detected with the two Tspan HiBiT sensors-loaded EVs (Figure 2E). By contrast, incubation with EVs loaded with LacC2 sensor led to significant complementation, revealing cytosolic delivery of the EV content in the target cells (Figure 2E). This difference between Tspan and LacC2 sensors could not be attributed to the labelling of distinct EV populations by the respective sensors as approximately 70% of LacC2-HiBiT-positive EVs were also positive for at least one of these tetraspanins (Supplementary Figure 2B). We thus chose the LacC2 sensor to further challenge and characterize the mechanisms of EV content cytosolic delivery.

### Uptake and cytosolic delivery of EV-loaded HiBiT-LacC2

To estimate EV uptake, we compared the luminescence activity in lysed cells at 37°C to the one measured at 4°C, a temperature at which uptake, but not EV adsorption to the cell surface or substrate, is inhibited. Following the same reasoning, the 4°C condition was used as the measure of residual unspecific activity in cytosolic delivery settings. We named this extended protocol “LUCID-EV”, standing for LUCiferase-based Internalization and Delivery of Extracellular Vesicles. We first performed a kinetic analysis. Cytosolic delivery was detected as soon as 15 minutes after EV addition and reached a plateau at 30 minutes to 1 hour before decreasing (Figure 3A). This late decrease was unlikely due to EV loss in the medium as 60% of HiBiT-loaded intact EVs were still detectable by luminescence in the medium after 2 hours of incubation (Supplementary Figure 4A). Within the same time frame, cytosolic delivery measurement remained close to background values at 4°C (Figure 3A). The kinetics of uptake (calculated as the difference between 37°C and 4°C measurements, Supplementary Figure 4B) and cytosolic delivery were similar up to 1 hour (Figure 3A, B), but uptake (and not cytosolic delivery) further increased at 2 hours before going down at 4 hours (Figure 3B, Supplementary Figure 4B). Although measurable, EV cytosolic delivery was poorly effective relative to the initial EV input (0.01%). However, the efficiency of EV uptake showed a similar trend, never reaching more than 0.1% of the input.

**Figure 3:**
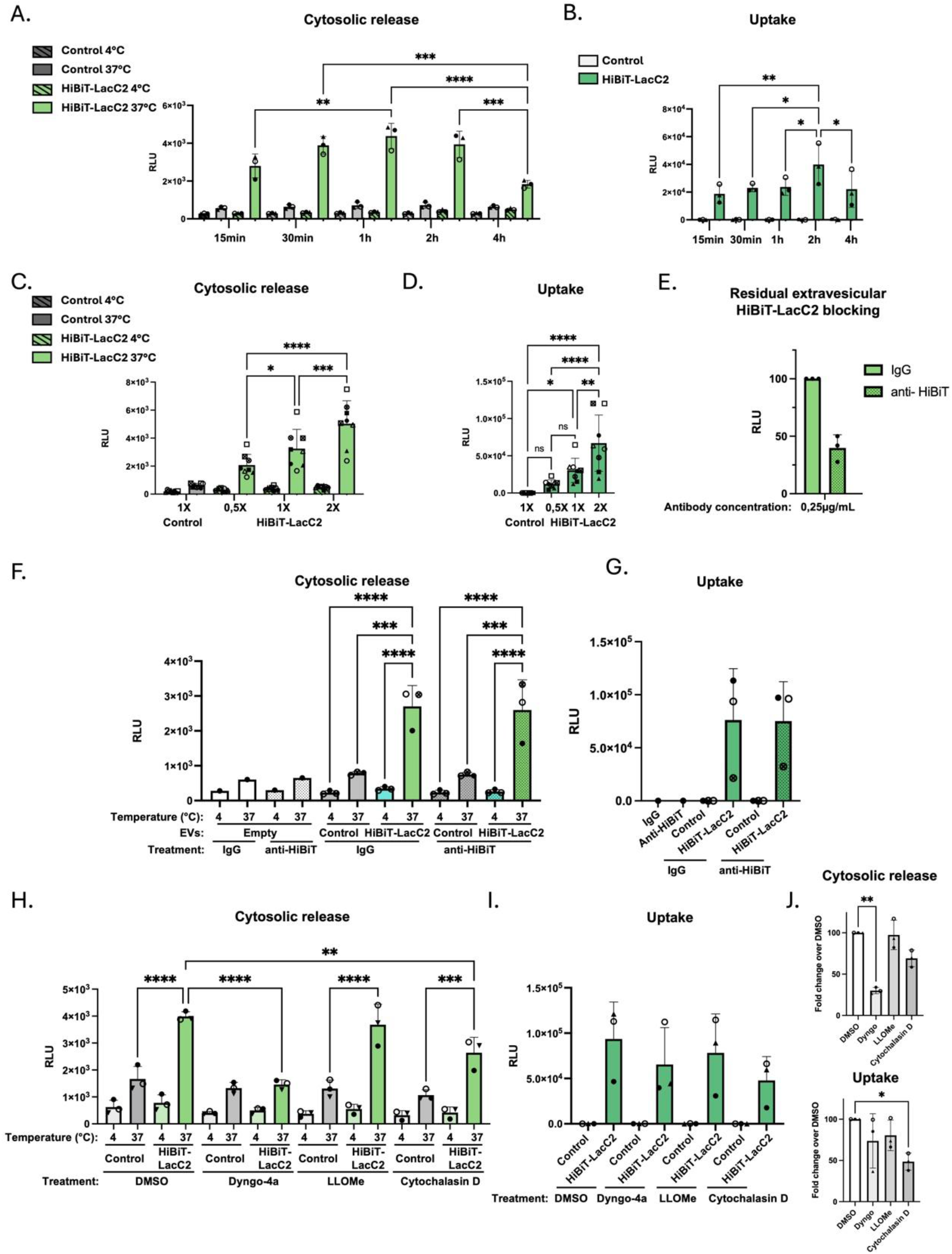
Time and dose-dependent EV cytosolic delivery. **A.** Time and temperature-dependent cytosolic release, and **B.** uptake of HiBiT-LacC2 or control EVs in MutuDC reporter cells. N=4 independent experiments. **C.** Temperature and dose-dependent cytosolic release and **D.** uptake of HiBiT-LacC2 or control EVs in MutuDC reporter cells for 2h. N=8 independent experiments. N=8 independent experiments. **E.** HiBiT-LacC2 detected on intact EV surface pre incubated with anti-HiBiT or IgG measure by bi-molecular complementation. N=3 independent experiments **F.** Cytosolic release, and **G.** uptake of EVs pre-treated with IgG or anti-HiBiT for 2h in MutuDC mpLgBiT. N=3 independent experiments. **H.**Cytosolic release, and **I.** uptake of HiBiT-LacC2 or control EVs incubated 30minutes with MutuDC mp-LgbBiT reporter cells pretreated for 45minutes with the following inhibitors: 12,5µM dyngo-4a (dynamin inhibitor), 0,3mM L-Leucyl-L-Leucine methyl ester hydrobromide (LLOMe) (endolysosomal disruptor), 10µM Cytochalasin D (actin polymerization inhibitor). **J.** Fold change of drugs to DMSO representation for HiBiT-LacC2 cytosolic delivery (based on Figures 3H) and uptake (based on figure 3I) Mean ± SD. Statistical analyses were performed using repeated measures non parametric one-way anova with Bonferroni test for A-J panels. \**P* < 0.05, \*\**P* < 0.01, \*\*\**P* < 0.001 and \*\*\*\**P* < 0.0001.

Since EV cytosolic delivery seems saturable at late time points, we performed dose response experiments, using three different amounts of EVs, ranging from 0.5 to 2X of the EV concentration used previously (i.e. from 0.5×10^7^ to 2×10^7^ RLU EVs, equivalent to1.25×10^8^ to 5×10^8^ particles). After 2 hours of incubation, we observed a robust dose-dependent increase in both EV uptake and cytosolic delivery (Figure 3C,D, Supplementary Figure 4C). It indicates that neither cytosolic delivery nor uptake are saturable within the tested ranges, suggesting that efficient EV interaction with the plasma membrane might be a limiting step in these processes. We also assessed the contribution of the surface-associated (non-luminal) pool of HiBiT-LacC2 (Figure 1E, Figure 3E-G). Incubation of intact HiBiT-LacC2 EVs with an anti-HiBiT antibody resulted in an almost twofold reduction of the luminescence signal after incubation with recombinant LgBiT protein (Figure 3E). This pretreatment affected neither EV uptake nor cytosolic cargo release of HiBiT-LacC2 EVs despite masking the EV surface-associated pool (Figure 3F,G), indicating a negligible contribution of extravesicular HiBiT to the measure of both mechanisms.

To further investigate the mechanisms involved in EV uptake and cytosolic delivery, recipient cells were treated with a panel of pharmacological modulators prior to EV incubation. Given the well-established phagocytic and macropinocytic activity of dendritic cells, we used cytochalasin D to inhibit actin polymerization, thereby impairing phagocytosis and macropinocytosis. In addition, we used Dyngo-4a, a dynamin inhibitor which blocks membrane scission, and L-leucyl-L-leucine methyl ester (LLOMe), which induces endo-lysosomal membrane disruption (Figure 3H-J). Dyngo-4a markedly decreased cytosolic delivery while having little effect on EV uptake. Cytochalasin D induced a modest reduction in both EV uptake and cytosolic delivery. In contrast, LLOMe had only limited effects on both processes under the same conditions (Figure 3H-J).

From these results, we conclude that the LUCID-EV assay allows quantitative and sensitive measurement of both EV uptake and cytosolic delivery. Importantly, the assay can be combined with pharmacological perturbations to investigate the cellular mechanisms governing these processes. In the tested system, cytosolic delivery is more sensitive to dynamin inhibition than EV uptake and suggest that actin-dependent uptake pathways contribute, at least in part, to both EV internalization and subsequent cytosolic delivery.

### Cytosolic delivery of EVs from different cell lines

The nature of EV-producing and of EV-recipient cells likely modulates the efficiency of EV uptake. We thus compared the fate of EVs produced by different cell lines but incubated with the same MutuDC reporter cell line. Similarly to E0771 cells, we engineered MutuDC, human mammary breast cancer MDA-MB-231 and human embryonic kidney HEK-293 cell lines to express HiBiT-LacC2. Normalised by the number of particles, similar efficiency of HiBiT-LacC2 loading into EVs was observed except for HEK 293 cells with an almost 10-fold increase (Figure 4A), mirroring the levels of expression of the sensor in cell extracts (Supplementary Figure 5A). The percentages of HiBiT-positive EVs produced by the different cell lines, as measured with Nano flow cytometry, were similar, ranging from 50% to 80%. However, we also noticed an unexpected staining of up to 40% of intact HEK-293 EVs (Figure 4B), suggesting reduced EV integrity or stability, or unusual LacC2 targeting at the EV surface with this cell line. As shown above for E0771 HiBiT-LacC2 EVs (Figure 3E-G), the relatively abundant surface-exposed HiBiT in HEK-293 EVs likely did not contribute to the detected cytosolic delivery signal, since, after 2 hours of incubation with MutuDC reporter cells, the different EV types all displayed similar dose-dependent EV cytosolic delivery, uptake and delivery efficiency (Figure 4C-F). These results demonstrate that the LUCID-EV assay could be applied to different cell contexts, at least regarding the source of EVs.

**Figure 4:**
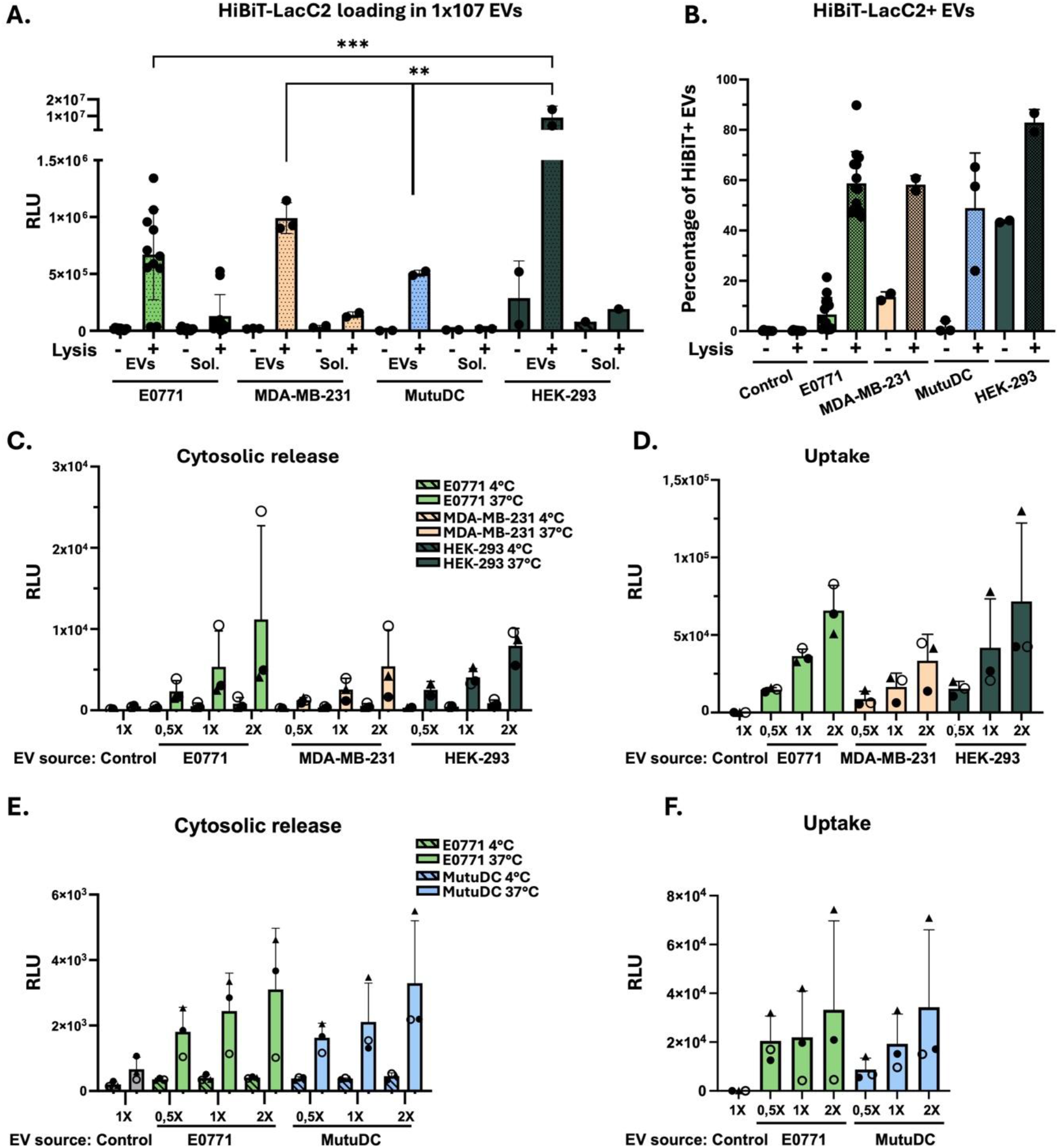
Capture, uptake and cytosolic delivery of HiBiT-LacC2 EVs from different secreting cells. **A.** HiBiT measurement by complementation assay with synthetic LgBiT protein in 10^6^ EVs from the indicated cell lines. N= 9 for E0771, N=3 for MDA-MB-231, N= 2 for MutuDC, N=2 for HEK. **B** Immunodetection by nano-flow cytometry of HiBiT-LacC2 positive EVs from the indicated cell lines lines, after or not saponin permeabilization (Lysis +/-). N=9 for E0771 EVs, N= 2 for MDA-MB-231, and HEK-293; N=3 for MutuDC, measured **C.** Cytosolic release, and **D**. uptake of HiBiT-LacC2 or control EVs from the indicated cell lines in MutuDC reporter cells after 2 hours of incubation. N=3 independent experiments. **F.** Cytosolic release, and **G**. uptake of HiBiT-LacC2 or control EVs secreted from the indicated cell lines in MutuDC reporter cells after 2 hours of incubation. N=3 independent experiments. Statistical analyses were performed using one-way anova with Sidak’s multiple comparison for A-F panels. Error bars = Mean ± SD. \**P* < 0.05, \*\**P* < 0.01, \*\*\**P* < 0.001 and \*\*\*\**P* < 0.0001.

### VLPs and small EVs deliver their cargo in MutuDC

Multiple subtypes of EVs are secreted concomitantly by a single cell type. In E0771 cells, as in other mouse cancer cell lines, we previously showed that a specific set of particles called viral-like particles (VLPs) is secreted due to the expression of endogenous retroviral genes (Cocozza et al., 2023). We also previously established that VLPs can be separated from EVs with a velocity density gradient (Figure 5A) (Cocozza et al., 2023). We asked whether VLPs and EVs behave similarly in the LUCID-EV assay. After density gradient separation, total protein staining revealed distinct protein profiles in fractions 1-3 and 4-6 (Figure 5B), in agreement with our previous study (Cocozza et al., 2023). Focusing on the most distinct fractions (fractions 1-2 and 5-6), Cd9 and Cd63 EV markers were enriched in low-density fractions (fractions1-2; Figure 5C) and the endogenous retroviral gag protein in VLP fractions (fractions 5-6; Figure 5C). HiBiT-LacC2 was detected in all fractions to a similar extent relative to particle number (Figure 5C-D). Each of the 4 fractions was evaluated separately, as their size and density profile differ from one another (Figure 5C,E). After three hours of incubation with MutuDC reporter cells, both HiBiT-LacC2-tagged EVs and VLPs were capable of cytosolic delivery (Figure 5F) and uptake (Figure 5G) with similar efficiencies.

**Figure 5:**
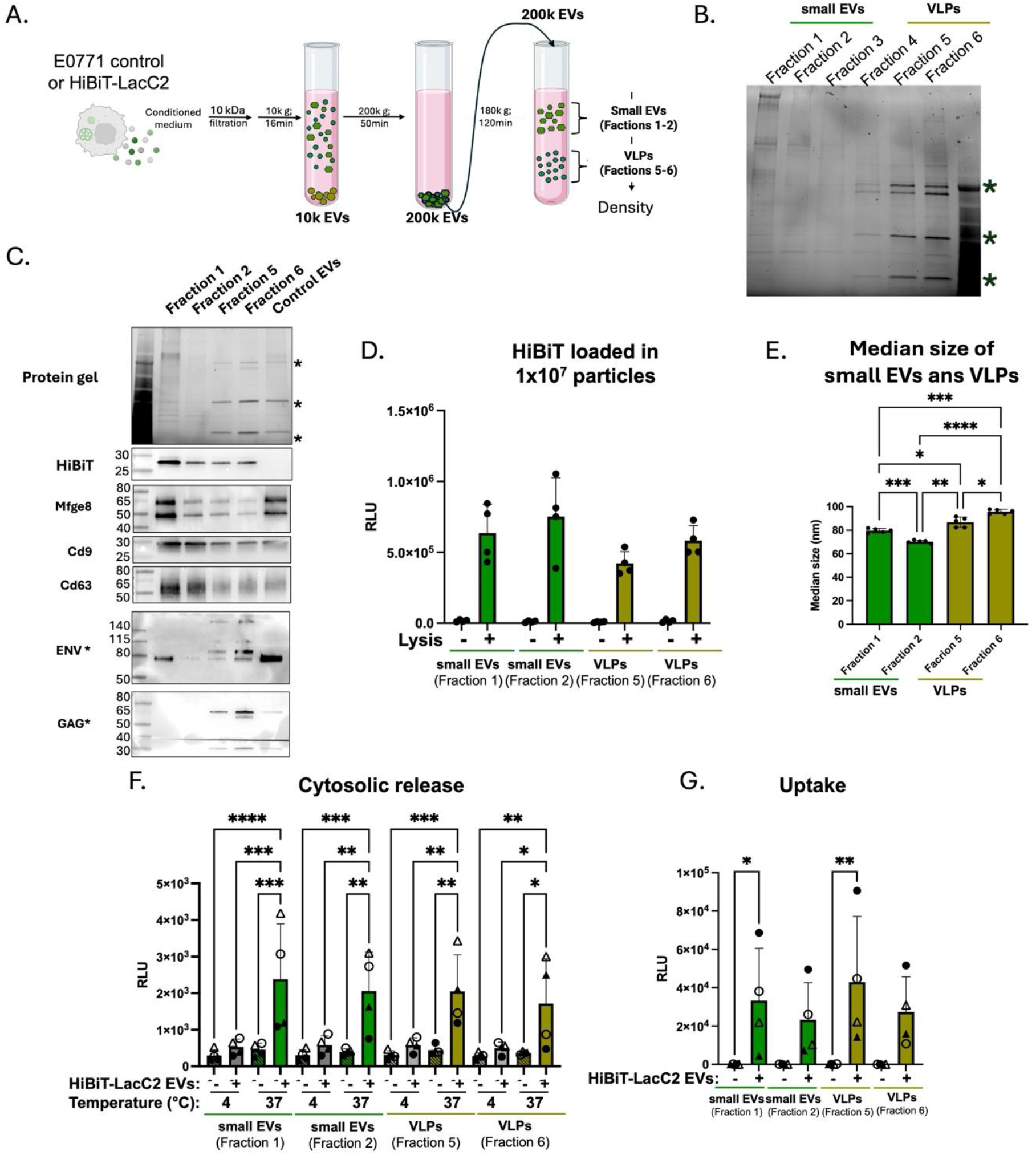
Uptake and cytosolic delivery of HiBiT-LacC2 small EVs and VLPs. **A.** Protocol of small EVs and VLPs separation. Created with Biorender. **B.** Protein gel staining of small EVs and VLPs fractions. (*) viral proteins. **C**. Protein gel staining and western blot detection with the indicated markers of small EVs and VLPs fractions and SEC isolated control EVs (control EVs) (*) viral proteins. **D.** HiBiT measurement by complementation assay with recombinant LgBiT protein in 1×10^7^ particles. N=4 **E.** Median size distribution of particles from small EVs and VLPs fractions measured by nano-flow cytometry. **F.** Cytosolic release, and **G.** uptake of HiBiT-LacC2 or control small EVs or VLPs in MutuDC reporter cells after 3 hours of incubation. N=4. Statistical analyses were performed using one-way ANOVA with Bonferroni test for E-G panels. Error bars = Mean with SD. \**P* < 0.05, \*\**P* < 0.01, \*\*\**P* < 0.001 and \*\*\*\**P* < 0.0001, not shown if not significant.

### Impact of EV purification on the detection of cytosolic delivery

EVs can be purified from the conditioned media of cell culture by multiple methods, each potentially resulting in the enrichment in specific EV subtypes or affecting EV composition, such as corona coating of the EV surface. We thus compared the fate of EVs either purified by SEC (usual protocol for our assay) or directly from the cell culture concentrated conditioned media (CCM), only depleted of cell debris and apoptotic bodies by low-speed centrifugation. Despite the presence of the bulk of soluble proteins in the CCM, HiBiT-LacC2 were mainly detected only after lysis treatment (Figure 6A), corroborating its enrichment in EVs. Cytosolic delivery from SEC-purified and from CCM-derived EVs were almost indistinguishable (Figure 6B). By contrast, the uptake of SEC-purified EVs was significantly higher than that of CCM, up to 3-fold (Figure 6C). The kinetics and dose-response of cytosolic delivery and uptake did not significantly differ between CCM and SEC-purified EVs (Figure 3A-D, Supplementary Figure 6A-D). We next assessed whether the addition of serum influenced EV uptake and cytosolic delivery (Figure 6 D,E). The addition of 10% EV-depleted serum to purified EVs (5% final) did not significantly affect either EV uptake or cytosolic delivery. These results further indicate applicability of the LUCID-EV assay using EVs isolated by different methods and in more physiologically relevant experimental settings.

**Figure 6:**
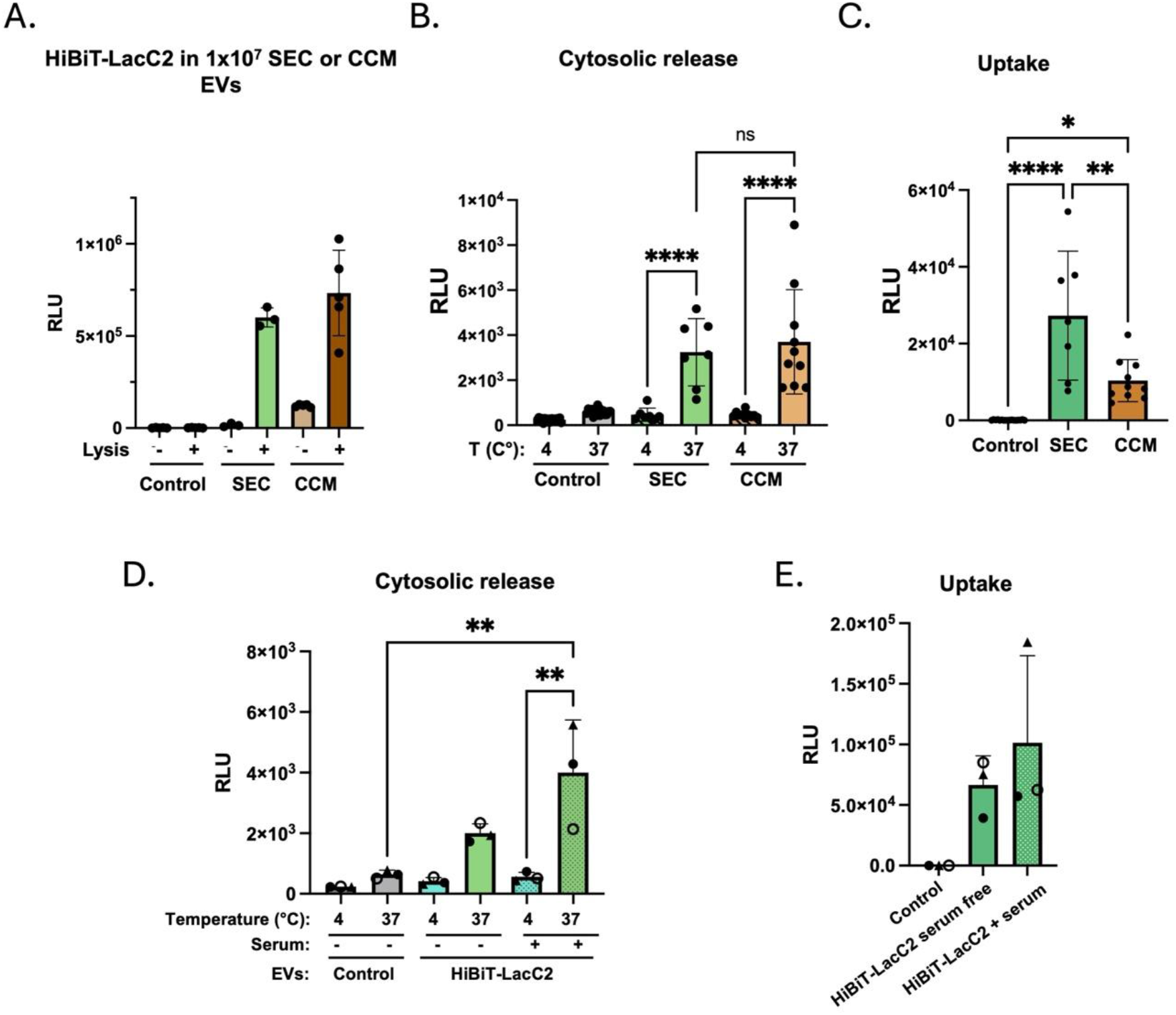
Influence of EV purification on EV cytosolic release efficiency. **A.** HiBiT measurement by complementation assay with recombinant LgBiT proteins in 1×10^7^ particles from SEC or CCM. N=3 for SEC EVs, N=5 for CCM **B.** Cytosolic release, and **C**. uptake of HiBiT-LacC2 or control CCM or SEC-purified EVs in MutuDC reporter cells. **D**. Cytosolic release, and **E.** uptake of control or HiBiT-LacC2 EVs incubated 2 hours with MutuDC mp-LgBiT reporter cells in serum free or 5% serum EV-depleted medium. N=3 independent experiments. Statistical analyses were performed using repeated measures non-parametric one-way anova with Bonferroni test for B-C panels. Error bars = Mean with SD. \**P* < 0.05, \*\**P* < 0.01, \*\*\**P* < 0.001 and \*\*\*\**P* < 0.0001. Comparison between HiBiT-LacC2 incubated at 37°C and controls (Control EVs at 4°c and 37°C; and HiBiT-LacC2 EVs at 4°C) are \*\*\*\**P* < 0.0001 but not shown for A-E panels. Statistical analyses were performed using T-test. Error bars = Mean with SD. \**P* < 0.05.

### Effect of VSV-G expression on EV cytosolic delivery

In most studies, detection of EV cytosolic delivery requires the presence of a fusogenic protein (usually VSV-G) at the EV membrane (Somiya and Kuroda, 2021; Liang et al., 2025; van den Ende et al., 2026). To evaluate the stimulatory effect of VSV-G expression in our assay, VSV-G protein was expressed by transient transfection in HiBiT-LacC2-transduced HEK-293 cells to produce EVs containing both proteins. VSV-G efficiently accumulated into EV, and its co-expression did not impair HiBiT-LacC2 EV-loading (Figure 7A, B), but the number of secreted particles was increased up to 10-fold by VSV-G expression (Figure 7C). After 30 minutes of incubation, EV cytosolic delivery was 20-fold higher in the presence of VSV-G (Figure 7D). The V-ATPase inhibitor bafilomycin A1 (BAF) is an efficient inhibitor of VSV-G fusogenic activity by blocking endosomal acidification. As expected, bafilomycin A1 treatment abolished VSV-G-induced cytosolic delivery. By contrast, it had little impact on delivery by EVs lacking VSV-G (Figure 7D). Uptake of VSV-G positive EVs was less affected by bafilomycin A1 treatment compared to cytosolic delivery (Figure 7E). The presence of VSV-G also significantly affected the kinetics of cytosolic delivery and uptake, showing a constant increase up to 4 hours (Figure 7F-G). Unexpectedly, the measure of VSV-G EV uptake, but not unmodified EV ones, was always lower than the measure of cytosolic delivery (Figure 7 F-G). It indicates that our protocol might underestimate the level of uptake, due to loss of material during the washing steps before cell lysis and to the limited amount of the cellular LgBiT pool released during lysis. We conclude that although our uptake assay protocol remains instructive for comparative analyses between conditions, it is less accurate for absolute quantification of VSV-G EV uptake specifically.

**Figure 7:**
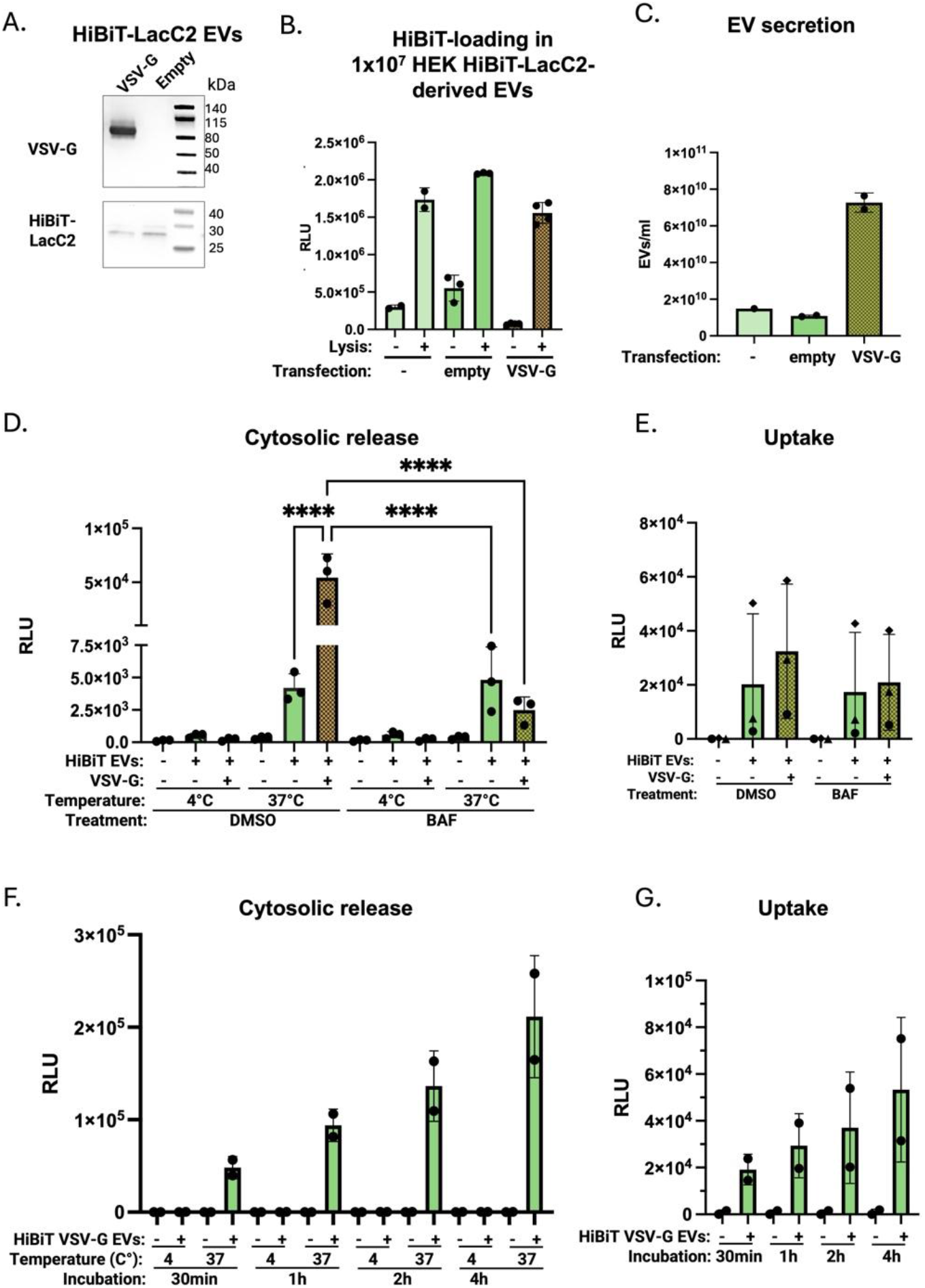
Effect of VSV-G expression on EV cytosolic delivery. **A.** Western blot of 5×10^8^ EVs derived from HiBiT-LacC2 expressing HEK-293 transfected with empty or VSV-G plasmid showing VSV-G (up) and HiBiT-LacC2 sensor (down). Representative of n=3 **B**. HiBiT measurement by complementation assay with synthetic LgBiT protein in intact or lysed HiBiT-LaC2 HEK-293 EVs transfected or not with empty or VSV-G plasmid. Normalized by 1×10^7^ particles. N=2 independent experiments for untransfected condition, N = 3 independent experiments for transfected condition. **C.** EV secretion normalized per the number of cells. **D.** Bafilomycin A1 (BAF) and temperature-dependent cytosolic delivery, and **E.** uptake of HiBiT-LacC2 (+/-), HiBiT-LacC2 + VSV-G (+/+) and control EVs (-/-) by DMSO or drug-treated MutuDC reporter cells after 30 minutes of incubation. N=3 independent experiments. **F.** Time and temperature-dependent cytosolic delivery, and **G**. uptake of HiBiT-LacC2-VSV-G (+) and control EVs (-) by MutuDC reporter cells, N=2 independent experiments. Statistical analyses were performed using one-way anova with Bonferonni multiple comparison for B-G panels, not shown if not significant. Error bars = Mean with SD. \**P* < 0.05, \*\**P* < 0.01, \*\*\**P* < 0.001 and \*\*\*\**P* < 0.0001.

### Role of LgBiT reporter intracellular localisation

HiBiT-LacC2 EVs significantly overlapped with HiBiT-pHluorin-Cd9 and HiBiT-pHluorin-Cd63 EV populations (Supplementary Figure 2B), indicating that the labelling of distinct EV subtypes could not account for the discrepancies observed in the measure of their cytosolic delivery. Since it was reported that EV cytosolic delivery preferentially occurs at the endosomal level (Joshi et al, 2020), we generated an additional MutuDC reporter cell line targeting LgBiT to the cytosolic face of endosomal membranes after fusion to a tandem PI3P lipid-binding FYVE domain derived from the ESCRT-0 protein Hrs (FYVE-LgBiT) (Figure 8A). As control, an untagged version of cytosolic LgBiT was also generated (Supplementary Figure 7A). The different reporter cell lines displayed variable levels of luminescence when assessed by direct complementation with an HiBiT peptide (Figure 8B, Supplementary Figure 7B). High expression of cytosolic LgBiT was associated with the release of LgBiT into the extracellular medium after 2 h of incubation (Figure 8C). Consequently, incubation of cytosolic LgBiT-expressing MutuDCs with control EVs resulted in a high background signal, which substantially reduced assay sensitivity and precluded reliable detection of EV-mediated cytosolic delivery in this reporter cell line (Supplementary Figure 7C-D). This problem was not observed with the FYVE-LgBiT reporter cell line which gave low level of cytosolic delivery signal upon exposure to control EVs at 37°C (Figure 8D). Strikingly, in contrast with mp-LgBiT, comparable levels of cargo delivery from HiBiT-pHluorin-Cd9, HiBiT-pHluorin-Cd63, and HiBiT-LacC2 EVs were detected in FYVE-LgBiT-expressing recipient cells (Figure 8D-E). These results suggest that the localisation of LgBiT in the reporter cell strongly impacts the measurement of EV cytosolic delivery.

**Figure 8:**
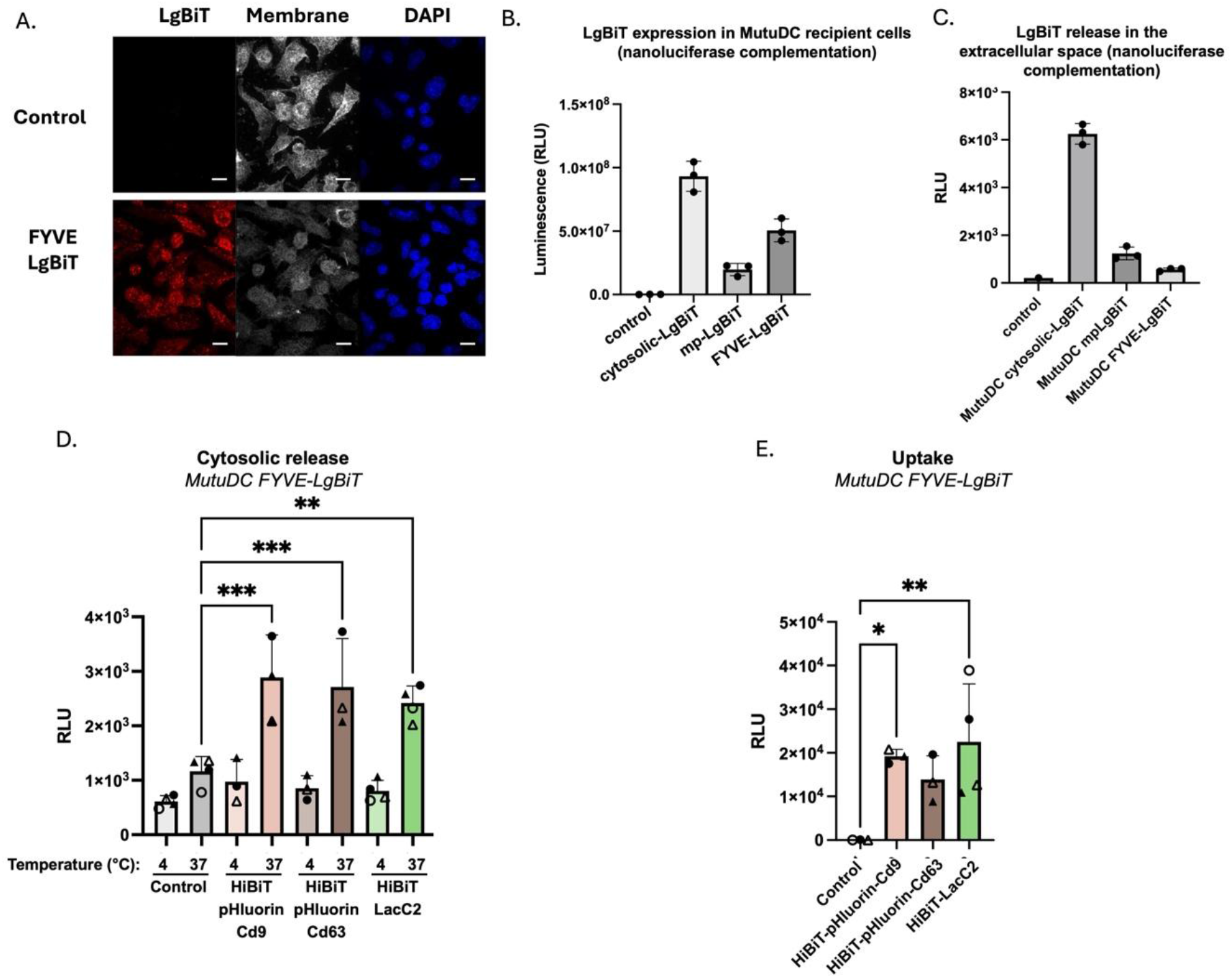
Role of LgBiT reporter intracellular localisation. **A.** Spinning disk microscopy images of MutuDC expressing or not FYVE-LgBiT, stained with anti-LgBiT (red) and WGA (plasma membrane, grey) and DAPI (nucleus, blue). Representative of N = 2. **B.** LgBiT expression in 150 000 MutuDC reporter cell lines (N=3 independent experiments) measured by complementation. **C.** Extracellular LgBiT released in the extracellular space by the different reporter cell lines measured by HiBiT complementation after 2h at 37°C. n=3 technical replicates **D.** Cytosolic delivery and **E.** uptake of HiBiT-pHluorin-Cd9, -pHluorin-Cd63 or -LacC2 EVs after 2 hours incubation with MutuDC FYVE-LgBiT, N=4 independent experiments

## Discussion

In this study, we report the design of an optimized assay based on the split HiBiT/LgBiT nanoluciferase enzyme, dedicated to the quantification of EV content cytosolic delivery in dendritic cells. The HiBiT/LgBiT assay offers a unique sensitivity suitable for the quantification of EV content cytosolic delivery, but it relies first on the efficient loading of HiBiT-tagged sensors into EVs and secondly on the complementation between the two fragments that, besides quantitative considerations, depends on their accessibility and respective localisations. Two alternative strategies, based respectively on the fusion of the HiBiT tag with Tspans or lipid binding domains were evaluated to promote EV loading. From the 7 tested sensors, 4 of them were poorly detected in cell extracts with the complementation assay, due to low expression and/or compromised interaction with the recombinant LgBiT fragment (Supplementary Figure 1 B-E). Insertion of a linker protein (pHLuorin) between the HiBiT tag and the Tspan CDS greatly enhanced luminescence detection (Supplementary Figure 1E). Three sensors, HiBiT-pHluorin-Tspans and HiBiT-LacC2, were efficiently loaded into EVs at similar levels (Figure 1C). Contrasting with Tspans constructs, Nano-flow cytometry identified surface-associated HiBiT-LacC2 in a minor fraction of EVs (6.6±6.6%), but we confirmed that most of HiBiT-LacC2 was protected from proteinase K degradation in intact EVs (Figure 1E, Supplementary Figure 2A).

Next, to optimise our assay, the LgBiT reporter expressed by the recipient cells was addressed to the cytosolic leaflet of the membrane compartment through addition of a post-translational myristoylation/palmitoylation sequence (mp) (Krager et al., 2012; Akdag et al., 2020), to promote its interaction with the membrane-bound EV sensors. Among the three tested sensors, only the LacC2 sensor was consistently detected into the cytosol following EV incubation with mp-LgBiT reporter cells (Figure 2E). Anti-HiBiT pre-treatment of HiBiT-LacC2 EVs efficiently reduced the detection of extravesicular HiBiT, but did not affect EV uptake or cytosolic delivery, thus excluding a main contribution of surface-accessible HiBiT-LacC2 in this process (Figure 3E-G). The substantial colocalization of LacC2 with Tspan markers detected by nano-flow cytometry in single EVs (Supplementary Figure 2B) makes unlikely that labelling of distinct EV subtypes by these two types of sensors could account for their different cytosolic delivery abilities. However, we cannot formally rule-out that the HibiT-LacC2 sensor is addressed to EVs presenting a specific lipid signature associated with enhanced cytosolic delivery.

Importantly, addressing LgBiT to the cytosolic face of endosomes upon fusion to a PI3P-binding FYVE domains in the recipient MutuDC cells allowed to detect the cytosolic delivery of the three HiBiT sensors with similar efficacies (Figure 8). By contrast, unmodified LgBiT reporter cell line was ineffective to detect cytosolic delivery with either of the three HiBiT sensors (Supplementary Figure 7C-D), but the higher activity (10 folds), but also background observed with this cell line in complementation assays (Figure 8B-C, Supplementary Figure 7B) and the non-canonical nuclear accumulation of the reporter (Supplementary Figure 7A) likely impair the detection of the process. These results demonstrate that in addition to the nature of the HiBiT sensor, the localisation of the LgBiT reporter is critical for the detection of EV cytosolic delivery. FYVE-LgBiT reporter concentrates at the cytosolic face of endosomes (Figure 8A), contrasting with the more diffuse distribution of mp-LgBiT, especially at the plasma membrane (Figure 2B). Although we can only speculate at this stage, mp- and FYVE-LgBiT reporters might detect two alternative cytosolic delivery routes occurring respectively at the plasma and endosomal membranes. The slow diffusion of the Tspan transmembrane sensors would restrict their interaction with LgBiT reporter at the plasma membrane but not in the restricted area of the endosomal membrane. More broadly, targeting LgBiT to distinct intracellular compartments could provide a strategy to investigate where along the intracellular trafficking pathway EV cargo becomes accessible to the cytosolic compartment.

In mp-LgBiT MutuDC recipient cells, the cytosolic delivery of HiBiT-LacC2 EV content is both time- and dose-dependent and inhibited at 4°C (Figure 3A). Relative to the starting EV input (10^7^ RLU), cytosolic delivery is poorly effective, reaching 0.01% at 2 hours of incubation (Figure 3B). Similarly, uptake does not exceed 0.1% of the starting input (Figure 3C). Importantly, this limited efficiency is consistent with previous studies using functional reporter systems, which have shown that only a small fraction of internalized EV cargo becomes functionally accessible in the cytosol, whereas a substantial proportion remains confined within endosomal compartments and may ultimately be degraded or re-released. Thus, low cytosolic delivery appears to represent a general bottleneck in EV-mediated cargo transfer rather than a limitation specific to our assay (Bonsergent et al., 2021 ; O’Brien et al., 2022 ; Van Solinge et al., 2023). Cytosolic delivery and uptake are not saturable in the range tested in our study as they increase concomitantly with the amount of EV input (Figure 3C,D). The kinetics of EV cytosolic delivery and uptake appear similar (Figure 3A) but these two processes are not strictly correlated. Indeed, when calculating the ratio of cytosolic delivery to uptake we observed that lower doses of EVs led to proportionally higher cytosolic delivery (Supplementary Figure 8A). Also, cytosolic delivery to uptake ratio of CCM-derived EVs is almost 4-fold higher compared to that of SEC-purified EVs (Supplementary Figure 8B). These results indicate that EV cytosolic release and uptake are not affected the same way by the dose and the EV isolation procedure.

Additionally, uptake is estimated by calculating the difference of luminescence activity in lysed cells (capture) at 37°C with the one measured at 4°C. At 4°C, EV absorption and non-specific EV binding is inhibited, but ligand-receptor interaction is taking place. Considering that EV uptake might be largely dependent on primary ligand-receptor interaction, our calculation is underestimating the uptake and the ratio of cytosolic release to uptake might be interpreted with caution (Supplementary Figure 8).

The four HiBiT-LacC2 EV preparations purified from different cell lines and species (mouse and human) showed similar cytosolic delivery and uptake efficacies (Figure 4). Similarly, EVs and VLPs behaved similarly in our assay (Figure 5). The apparent lack of specificity of EV cytosolic delivery according to the EV source suggests that we might detect only a basal cytosolic delivery activity with our set-up, potentially due to a non-optimal EV/reporter cell combination. Indeed, recent reports demonstrated that the efficiency of EV cytosolic delivery could involve specific ligand/receptor combination (Dancourt et al., 2026).

The ability to detect low-frequency delivery events is particularly important when investigating their potential biological relevance in physiologically meaningful recipient cells, including immune cells exposed to tumour-derived EVs. MutuDC cells were chosen in our assay because they are known functional target cells of E0771 EVs (Cocozza et al., 2023). Myeloid cells are also widely described in the literature as the main recipient cell type of EV uptake (Makhijani and McGaha, 2022; Lopera et al., 2026), making MutuDC a relevant physiological cellular model but with specific features compared to other cell types. They are specialized in phagocytosis and antigen cross-presentation. Cytochalasin D that inhibit both phagocytosis and macropinocytosis had only partial effect on EV cytosolic delivery. LLOMe, which stimulates antigen cross presentation (Gros et al., 2022), did not increase EV cytosolic delivery, indicating only a mild contribution of these pathways in EV cytosolic delivery in these cells. The fast kinetics of EV cytosolic delivery with mp-LgBiT recipient cells would fit with direct EV fusion events with the plasma membrane but conversely, Dingo-4a treatment, which is described to inhibit clathrin-mediated uptake, significantly inhibited EV cytosolic delivery (Figure 3H-J). The panel of compounds used in this study indicates a partial contribution of phagocytosis and macropinocytosis and a role of clathrin-mediated mechanisms in EV delivery, without allowing unambiguous distinction between plasma membrane fusion and endosomal entry routes. Further investigation using other reporters, such as the FYVE-LgBiT, and other recipient cell types will be required to further decipher delivery mechanisms.

Previous studies based on HiBiT/LgBiT split luciferase complementation concluded on the requirement of VSV-G expression on EVs to detect effective cytosolic delivery (Somiya and Kuroda, 2021). In our set-up, VSV-G expression increased up to 20 folds the efficacy of cytosolic delivery compared to EVs devoid of VSV-G, reaching 2.5% of the EV input after 4 hours of incubation (Figure 7F). The sensitivity of measurement thus appears to be the critical factor to detect cytosolic delivery. Contrasting with previous reports (Joshi et al., 2020; Bonsergent et al., 2021), EV cytosolic delivery is not affected by bafilomycin a1 treatment, which, as expected, fully inhibits VSV-G-induced EV fusion events.

In conclusion, LUCID-EV provides a sensitive and quantitative approach to monitor EV cytosolic delivery in living cells without artificial enhancement by fusogenic proteins. Our study demonstrates that EV-associated cargo can access the cytosol of dendritic cells under these conditions, while confirming that this process occurs at low efficiency. This low efficiency raises the question of the physiological significance of this process, which however could reflect suboptimal EV/recipient cell combination. Although our assay does not establish whether this level of cytosolic delivery is sufficient to induce a biological response, continuous EV uptake and processing under physiological conditions could allow low-frequency delivery events to accumulate over time and eventually produce biologically meaningful effects (O’Brien et al., 2022).Moreover, previous studies have demonstrated functional delivery of EV-encapsulated proteins and RNAs to recipient cells in vitro and in vivo, supporting the possibility that relatively infrequent cytosolic delivery events can contribute to EV-mediated intercellular communication (Zomer et al., 2015; de Jong et al., 2020; Ripoll et al., 2026). Taken together, LUCID-EV provides a framework to investigate the cellular mechanisms regulating this process and to further address the contribution of luminal cargo release to EV-mediated intercellular communication.

## Acknowledgments

We thank P. Panagiotis (Institut Curie, Paris) for generating the MDA-MB-231 HiBiT-LacC2 cells, and the CurieCoreTech EVs for help in use of the Flow NanoAnalyzer. This work was funded by INSERM, CNRS, Institut Curie, SIRIC Curie/INCa-DGOS-Inserm-ITMO Cancer_18000, PSL research University, BioSPC Doctoral School of Univ Paris Cité, program France 2030 launched by the French government, and grants from french ANR (ANR-22-CE18-0012, ANR-22-CE16-0033, ANR-10-IDEX-0001-02 PSL), INCa (INCa_16083, INCa_16735), Fondation ARC (PGA12021020003189_3588), Fondation Chercher et Trouver, and ITMO Cancer of Aviesan within the framework of the 2021-2030 Cancer Control Strategy, on funds administered by Inserm for purchase of the U30 Flow NanoAnalyzer.

---

**Supplementary Figure 1:**
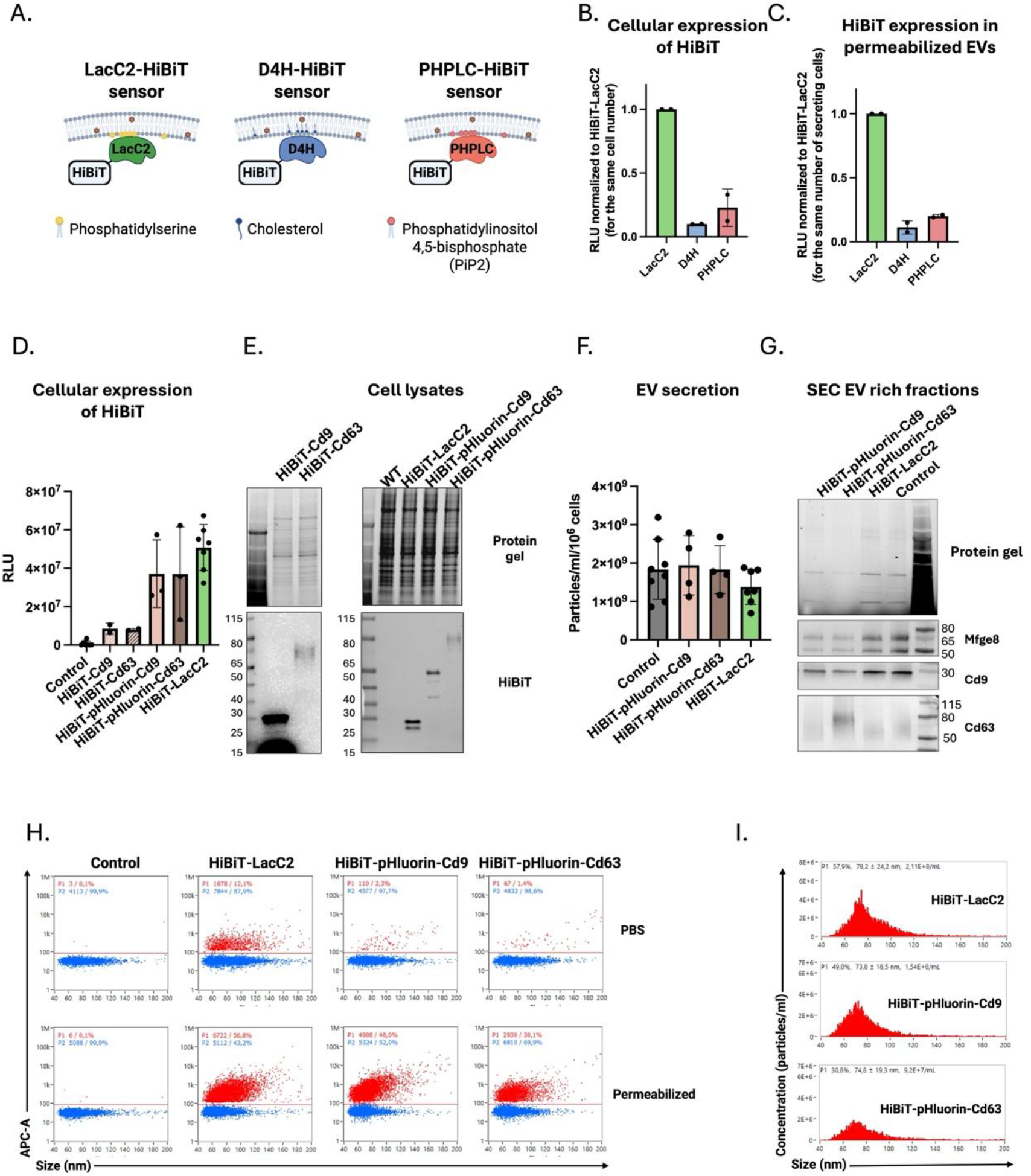
Strategies for HiBiT loading and characterization in EVs. **A.** Representation of the fusion protein with lipid sensors used to target HiBiT in EVs. Created with Biorender **B.** Measurement of Lipid-binding-HiBiT sensors by complementation assay with recombinant LgBiT protein in E0771 cell extracts, and **C.** permeabilized EVs. N= 2 independent experiments **D.** Western blot of E0771 expressing HiBiT fused to CD9, CD63, pHluorin-CD9, pHluorin-CD63 or C2 domain of lactadherin. Cell lysates from 100 000 cells showing total proteins (up) and HiBiT sensor(down). Representative of n=3 **E.** HiBiT measurement by complementation assay with synthetic LgBiT protein in 30 000 lysed E0771 cells. N= 2 for HiBiT-CD9, N=2 for HiBiT-CD63 N= 3 for HiBiT-pHluorin-CD9, N=3 for HiBiT-pHluorin-CD63, N=7 for HiBiT-LacC2. **F.** Western blot of 5×10^8^ EVs derived from control and HiBiT-LacC2 expressing E0771 cells showing CD81, AGO2, MFGE8, CD9 and CD63 proteins (from top to bottom) **G.** EV secretion normalized by 10^6^ secreting cells, measured by nanoFCM. Each dot represents a single independent experiment. **H.** Gating of HiBiT positive EVs under PBS and saponin treatment **I.** Histogram showing the size of HiBiT+ EVs (representative of N= 2 for HiBiT-pHluorin-CD9, N=2 for HiBiT-pHluorin-CD63, N=9 for HiBiT-LacC2 for panels C and D)

**Supplementary figure 2.**
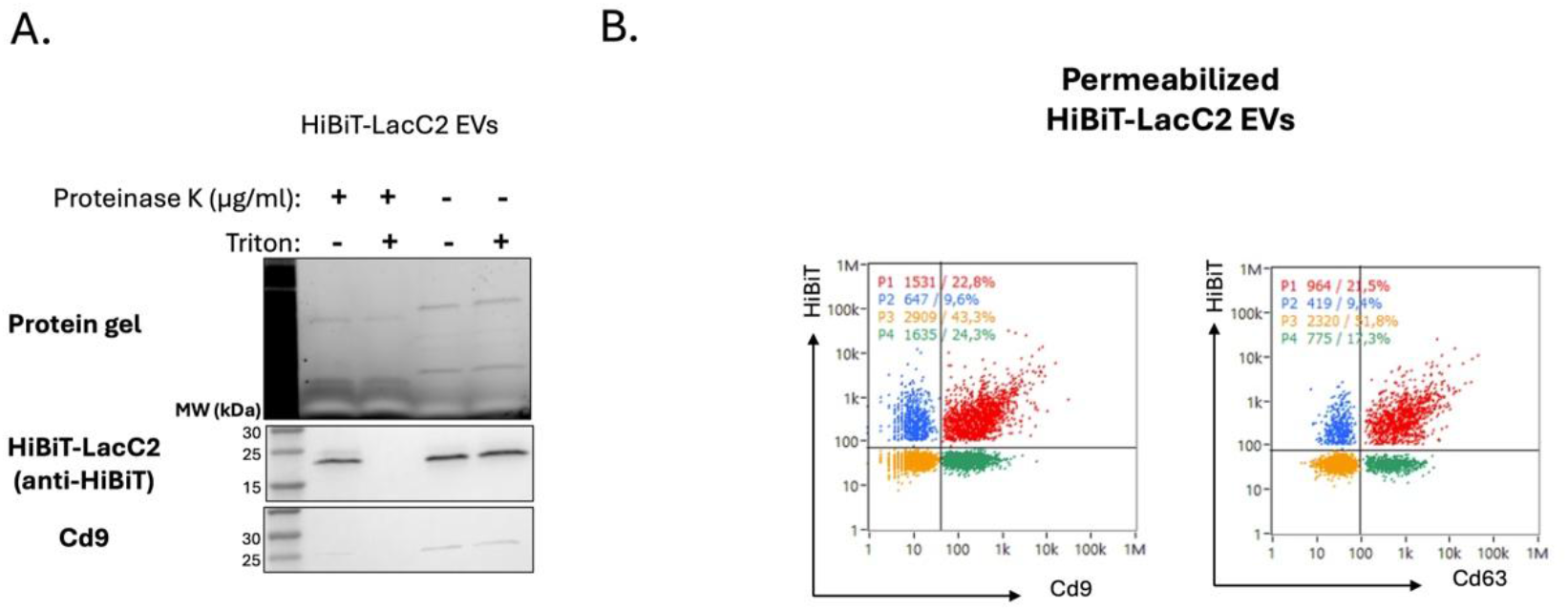
Characterization of HiBiT-LacC2 EVs. **A.** Western blot of SEC-purified EVs were treated with proteinase K in the absence or presence of 0.2% Triton X-100 showing protein gel, HiBiT-LacC2, MOV-10, and CD9 (from top to bottom). Representative of N=2 independent experiments. **B.** Permeabilized HiBiT-LacC2 EVs stained from CD9, CD63 and HiBiT staining analysed by nano flow cytometry

**Supplementary figure 3.**
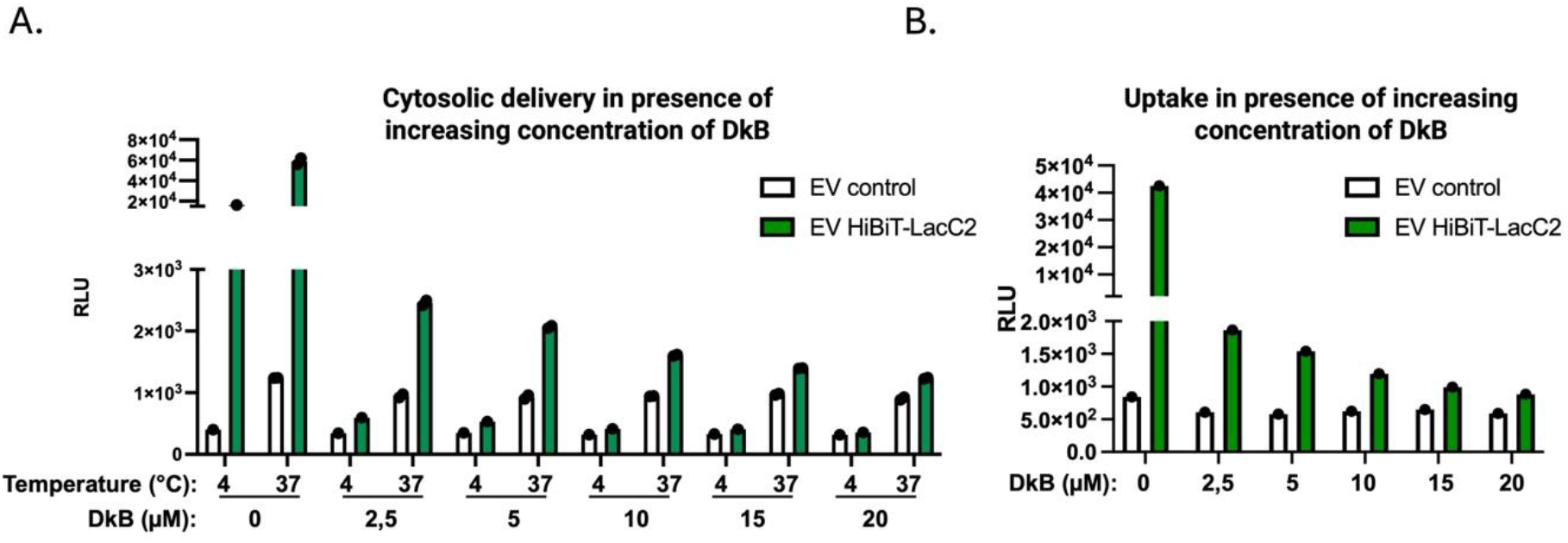
Titration of HiBiT EVs with soluble LgBiT and LgBiT cells with soluble HiBiT. **A.** Uptake and, **B.** cytosolic delivery of HiBiT-LacC2 or control EVs in response to increasing dose of DkB after 2 hours of incubation

**Supplementary Figure 4:**
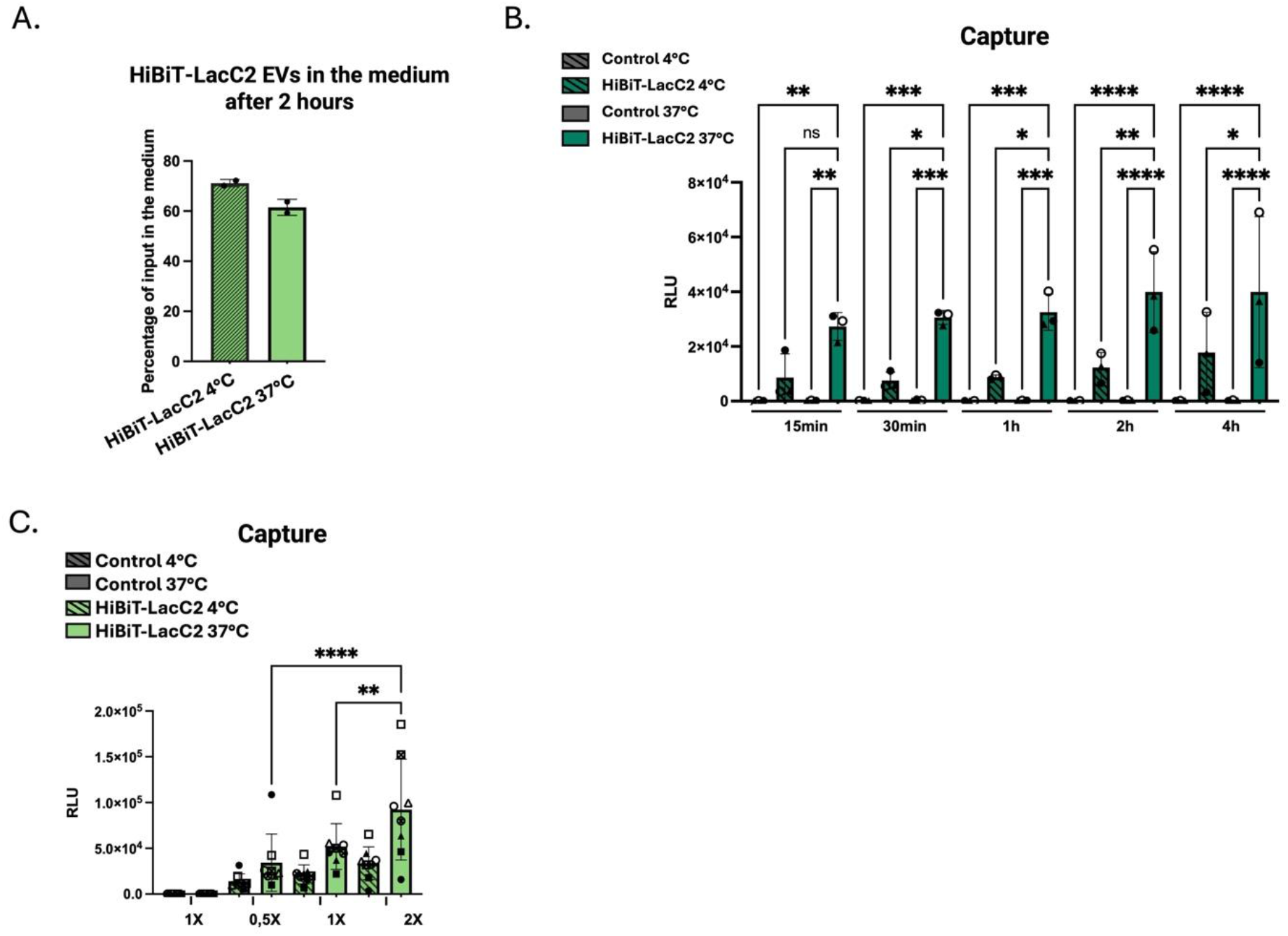
EV capture measured by LUCID-assay. **A.** Percentage of HibiT-LacC2 EVs left in the medium out of total input after 2 hours of incubation without DkB. N=2 technical replicates **B**. Time and temperature-dependent capture of HiBiT-LacC2 and control EVs by MutuDC expressing LCK-LgBiT. N=3 **C.** Temperature and dose-dependent capture of HiBiT-LacC2 and control EVs by MutuDC expressing LCK-LgBiT. N=8. Statistical analyses were performed using one-way anova with Bonferroni multiple comparison. Error bars = Mean with SD. \**P* < 0.05, \*\**P* < 0.01, \*\*\**P* < 0.001 and \*\*\*\**P* < 0.0001. Comparaison between HiBiT-C2 incubated at 37°C and controls (Control EVs at 4°c and 37°C; and HiBiT-C2 EVs at 4°C) are \*\*\*\**P* < 0.0001

**Supplementary Figure 5.**
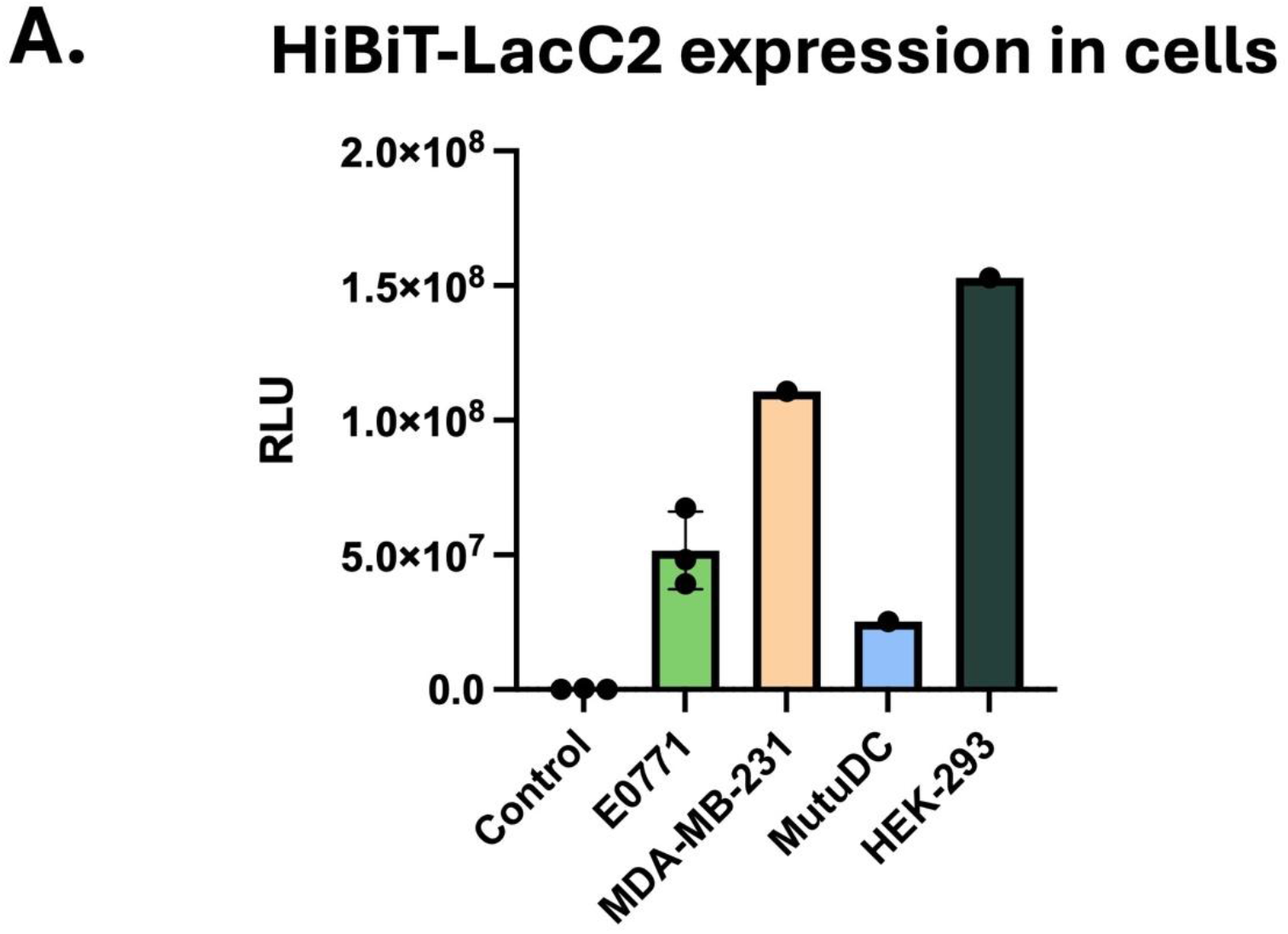
HiBiT-LacC2 expression in different secreting cell lines (change the title if we add capture data) **A.** HiBiT measurement by complementation assay with synthetic LgBiT protein in 30 000 lysed E0771 cells. N= 3 for E0771 control and E0771 HiBiT-LacC2, N=1 for MDA-MB-231, MutuDC and HEK HiBiT-LacC2. B.C.Capture?

**Supplementary Figure 6:**
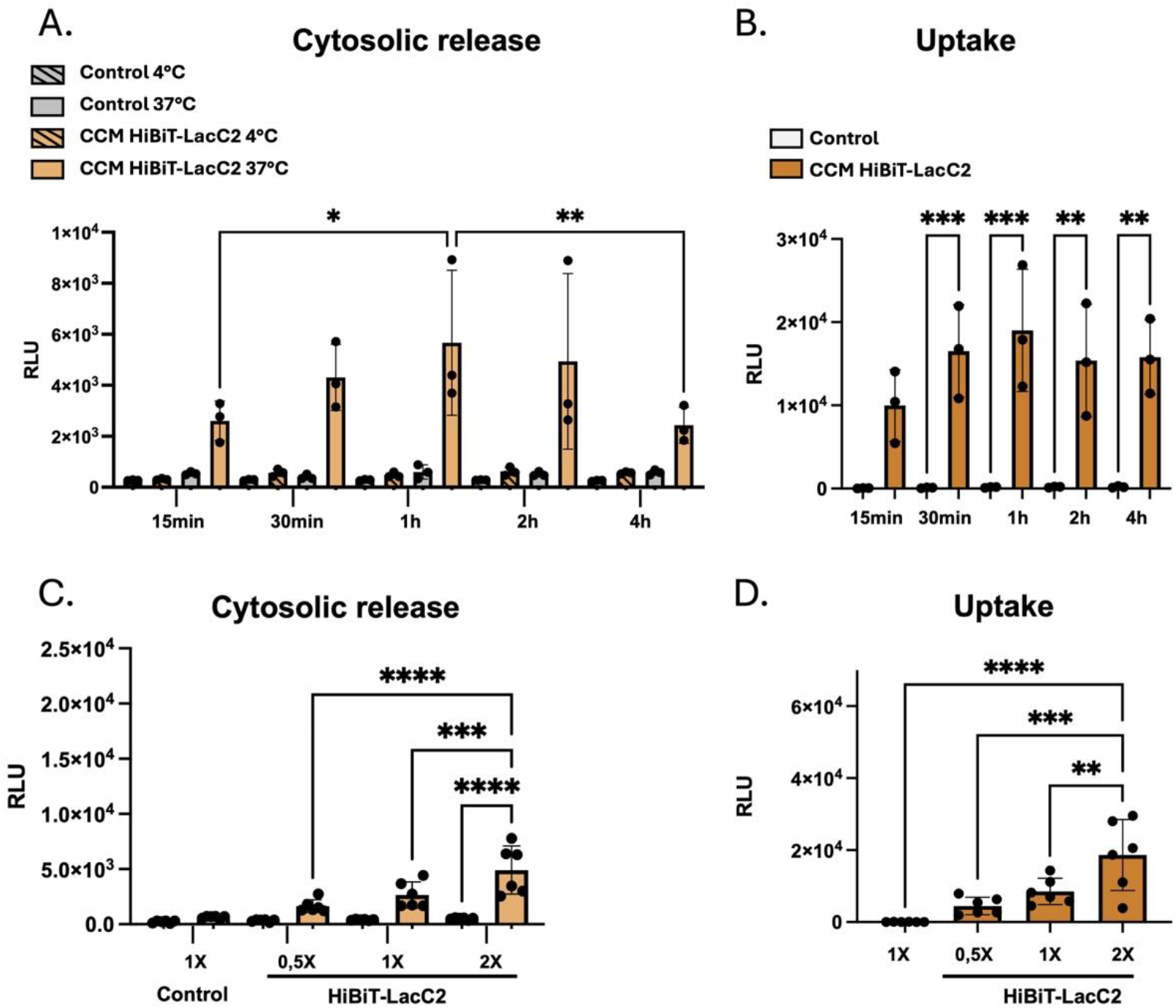
Uptake and cytosolic release HiBiT-LacC2 CCM EVs according to the time of incubation and EV dose. **A.** Time and temperature-dependent cytosolic release, and **B**. uptake of HiBiT-LacC2 and control EVs by MutuDC reporter cells. N=3 **C.** Temperature and dose-dependent cytosolic release, **D**. and uptake of HiBiT-LacC2 and control EVs by MutuDC reporter cells. N=6. Statistical analyses were performed using one-way anova with Bonferroni multiple comparison for A-F panels. Error bars = Mean with SD. \**P* < 0.05, \*\**P* < 0.01, \*\*\**P* < 0.001 and \*\*\*\**P* < 0.0001.

**Supplementary Figure 7:**
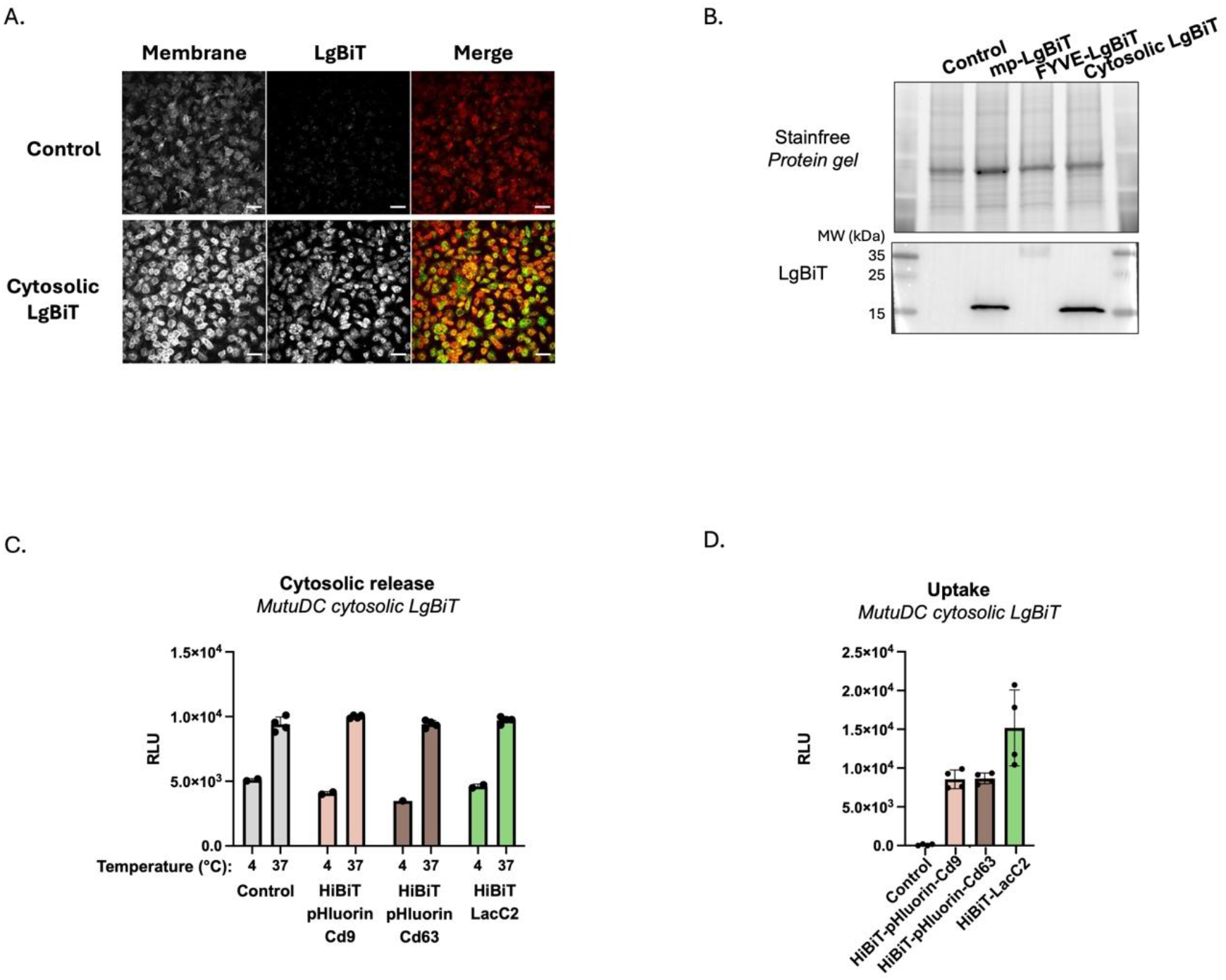
Characterization of MutuDC expressing reporter LgBiT in different subcellular localizations. **A.** Spinning disk microscopy images of MutuDC expressing or not mpLgBiT, stained with anti-LgBiT (green) and WGA (plasma membrane,red) and DAPI (nucleus,blue). Representative of N = 2 Scale bar = 10uM **B.** Western blot of cell lysate expressing the different LgBiT reporters showing the stainfree protein gel and the LgBiT reporter from top to bottom. **C.** Cytosolic delivery and **D.** uptake of HiBiT-LacC2 EVs after 2 hours incubation with MutuDC cytosolic LgBiT, n=4 technical replicates.

**Supplementary Figure 8:**
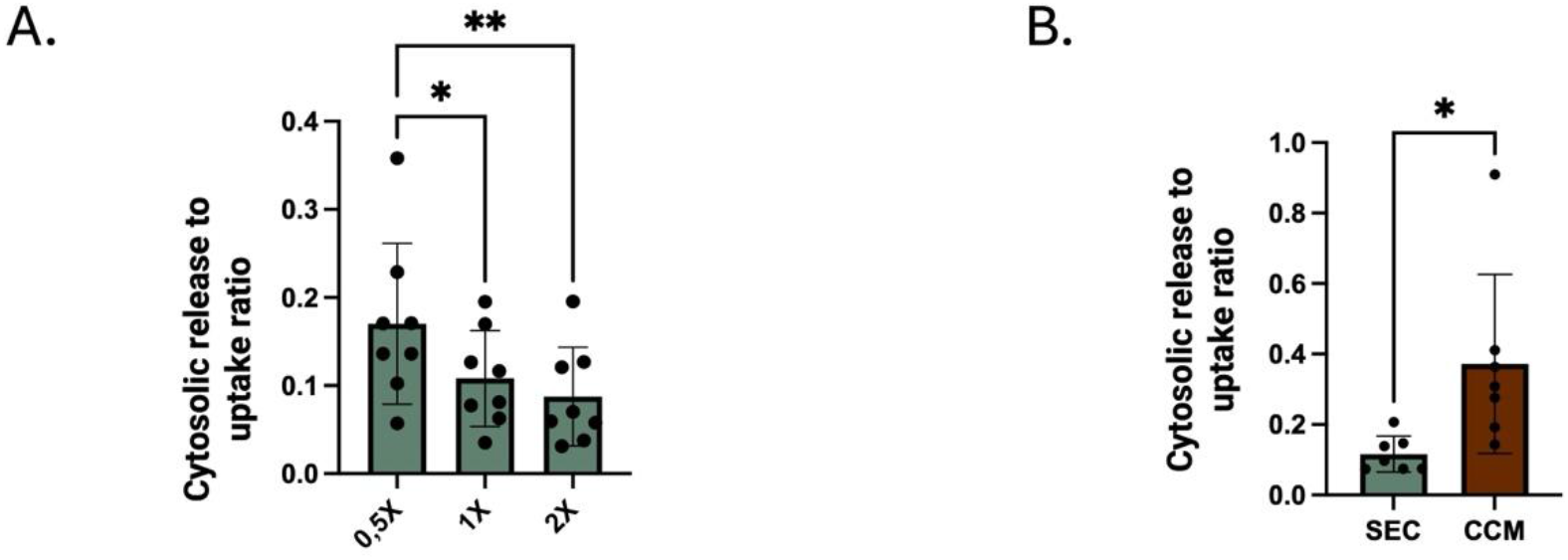
Cytosolic release/uptake ratio of HiBiT-LacC2 EVs in MutuDC reporter cells. **A.** Cytosolic release to uptake ratio of SEC purified HiBiT-LacC2 EVs based on Figure 3C-D **B**. Cytosolic release to uptake ratio. Error bars = Mean with SD. Data normalized to the 0,5×10^7^ RLU EVs condition based on Figure 6B-C. Statistical analyses were performed using one-way anova with Bonferroni multiple comparison for panel A and paired T test for panel B. Error bars = Mean with SD. \**P* < 0.05, and \*\**P* < 0.01.

